# The bacterial molecular switch DnaA-ATP integrates chromosome replication with cell growth and division dynamics

**DOI:** 10.1101/2025.02.22.639668

**Authors:** Ezza Khan, Paola E. Mera

## Abstract

Successful proliferation of bacteria requires the coordination of multiple cellular events. The precise coordination between the replication of the chromosome with cell growth and division is essential for daughter cells to inherit an intact copy of the genome with the appropriate cell size. However, the molecular mechanisms that temporally and spatially integrate these essential processes have remained elusive in the field. Using the model system *C. crescentus*, we uncovered a previously unrecognized role for the conserved chromosome replication initiator DnaA in regulating cell growth and division dynamics. Through mutant analyses, transcriptional profiling, biochemical and high-resolution imaging, our data demonstrate that DnaA modulates cell size through direct transcriptional activation of enzymes involved in cell wall biosynthesis. We identify a conserved cell wall biosynthetic enzyme, MurD, as a key downstream effector and demonstrate its unexpected ability to restrict cell size. Both *in vivo* and *in vitro* analyses revealed that the transcriptional regulation of *murD* requires the ATP-bound form of DnaA, the same nucleotide-bound form required for initiating chromosome replication. These findings highlight how bacteria leverage the conserved nucleotide-dependent switch, DnaA-ATP, to temporally orchestrate spatially distinct processes ensuring accurate progression of the cell cycle.

## INTRODUCTION

Bacteria must tightly coordinate multiple cellular processes to ensure successful progression of the cell cycle. This coordination involves the regulation, temporally and spatially, of key events like chromosome replication and segregation, as well as the elongation and constriction of the cell envelope^1^. A critical component of this regulatory circuitry includes the biosynthesis and maintenance of precursors that are necessary to sustain the forward progression of the cell cycle. In addition to the intracellular multi-level coordination, bacteria actively integrate environmental cues into the cell cycle regulatory circuitry to quickly adapt to changing conditions ^2^. How bacteria manage to coordinate all these processes while growing and dividing at such rapid rates remains a fascinating challenge for scientists to unravel. In this study, we aimed at uncovering molecular details of such strategies.

One of the first committed steps in the bacterial cell cycle is the initiation of chromosome replication, a process regulated by the conserved initiator protein DnaA ^3–5^. DnaA bound to ATP (DnaA-ATP) oligomerizes at the origin of replication (*oriC*), opens *oriC,* and allows the replication machinery to assemble and initiate chromosome replication ^3,4^. Following initiation of replication, the protein Hda bound to the β-clamp of the replisome turns off DnaA by promoting DnaA’s intrinsic ATPase activity ^6,7^. Besides its role in chromosome replication, DnaA serves as a conserved transcription factor ^8,9^, albeit fewer mechanistic details are known about this function. The DnaA transcriptional regulon commonly comprises genes encoding components of the replisome and enzymes involved in nucleotide biosynthesis ^9,10,11^. Interestingly, both functions of DnaA (replication initiation and transcriptional regulation) are modulated by DnaA’s nucleotide-bound state: ATP binding and hydrolysis are essential for its activity as a replication initiator^12,13^, while the identity of the bound nucleotide (ATP vs. ADP) can influence its function as a transcription factor^14–16^.

The ability of bacteria to double in size and undergo division relies on the coordinated assembly and remodeling of their cell wall ^17^. The cell wall is primarily composed of peptidoglycan (PG), a polymer of glycan strands consisting of alternating N-acetylglucosamine and N-acetylmuramic acid (GlcNAc & MurNAc) residues, cross-linked by short peptides ^18^. PG biosynthesis begins in the cytoplasm with the production of the nucleotide-linked precursors UDP-GlcNAc and UDP-MurNAc, which are subsequently assembled into lipid II by membrane-associated enzymes, and ends in the periplasm where protein complexes coordinate cell wall remodeling during growth and division ^19,20,21^. In rod-shaped bacteria, new cell wall material is typically inserted laterally during cell elongation by the Rod complex, which includes the actin homolog MreB ^22,23^. Subsequently, PG synthesis shifts to the midcell driven by the divisome machinery, which includes the tubulin homolog FtsZ ^24^. The molecular mechanisms that temporally integrate chromosome replication initiation with cellular expansion and division remain poorly understood.

Identifying the molecular mechanisms that coordinate multiple events over the progression of the cell cycle is inherently challenging due to the intricate and overlapping nature of these processes. However, the diderm oligotrophic bacterium *Caulobacter crescentus* offers a powerful model system that enables high-resolution analysis of cell cycle regulation due to its well-defined developmental stages and robust genetic tools^25–29^. Upon cell division, *C. crescentus* produces two genetically identical but morphologically distinct cells: a smaller motile swarmer cell that cannot initiate chromosome replication and a sessile stalk cell that is replication competent^29^ (Figure 1A). The dimorphic life cycle of *C. crescentus* enables isolation of homogeneous populations of cells at different developmental stages ^30–33^. Furthermore, *C. crescentus* initiates chromosome replication once per cell cycle regardless of nutrient conditions ^34^ enabling detailed investigation into the mechanisms that coordinate onset of chromosome replication with cellular growth and the progression of the cell cycle.

**Figure 1.**
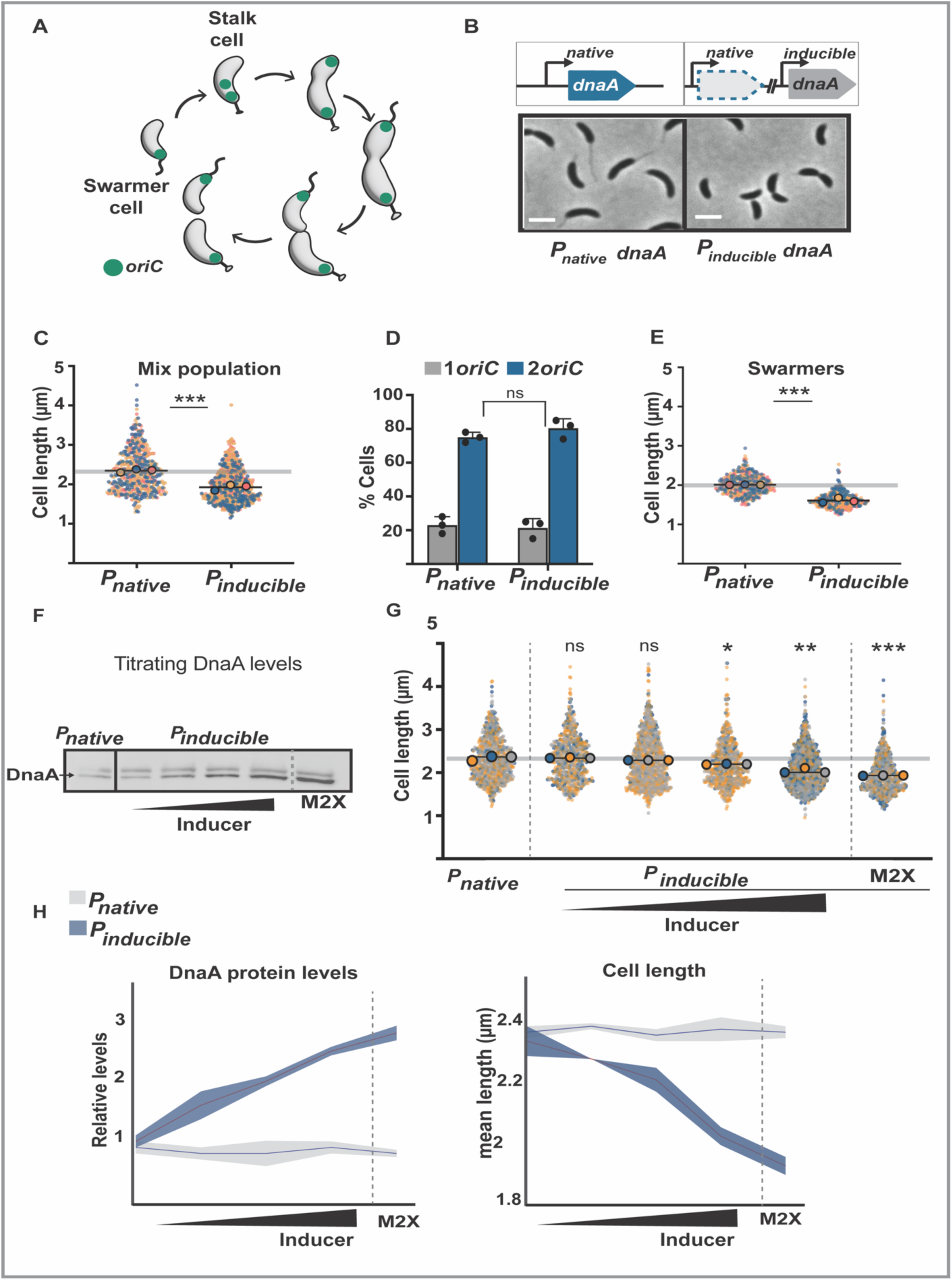
DnaA impacts cell size without changing *oriC* copy number. **A.** (Left) Schematic representation of *Caulobacter* dimorphic lifestyle. **B**. Phase contrast images of WT (*parB*::CFP-*parB*) and *dnaA* inducible strain (*dnaA*::Ω, *parB::cfp-parB, P_van_ dnaA)* grown in minimal media (M2G) to exponential phase, scale 2µm. **C.** Super plots showing cell length analysis of mixed population. Small dots represent data points from three independent replicates, large dots represent median values (blue, pink, yellow). The horizontal line represents the mean of three median values. *dnaA* inducible shows ∼20% decrease in cell length. **D.** Percentage cells representing *oriC* quantification in WT and inducible *dnaA* strain showing non-significant change. **E**. Super plots showing cell length analysis of synchronized population grown in M2G. *dnaA* inducible swarmer’s show 20% decrease in cell length. **F.** Western Blot showing DnaA titration using WT and inducible *dnaA* (*dnaA*::Ω, *parB::cfp-parB, P_xyl_ dnaA*). Cells were grown in M2G with xylose titrated from 0.025%, 0.05%, 0.075% to 0.3% in M2G, to exponential phase. For maximum expression cells were grown in M2X to shown after dotted line. **G.** Super plots showing cell length analysis of DnaA titration. Small dots represent data points from three independent replicates, large dots represent median values (blue,grey,orange). At 0.025% xylose cell length is comparable to WT, M2X shows decrease in cell length ∼20%. **H.** Quantification of DnaA levels left panel, DnaA levels increase with higher inducer percentage in comparison to WT in minimal media. Right panel shows quantification of cell length in comparison to WT; the maximum drop is observed with M2X. Data points show mean ± SD. A parametric t-test was performed using population mean values (N = 3) to compare values for each measurement. p(<0.0001) ****, p(0.0001)***, p(<0.05)*, ns (non-significant). All samples were blinded n= ∼600 cells.

A recurring theme in biology is that nature often achieves remarkable complexity through elegantly simple regulatory mechanisms. In this study, we uncovered one such mechanism in which the conserved initiator of chromosome replication DnaA regulates growth and constriction rate through the transcriptional regulation of cell wall biosynthetic enzymes. Specifically, we identify MurD, a key enzyme in the conserved cytoplasmic pathway for peptidoglycan precursor biosynthesis, as a direct target of DnaA. Our findings reveal that MurD plays an unexpected role in the regulation of cell size. Collectively, this work demonstrates a direct regulatory axis linking chromosomal maintenance to cell growth and size, positioning the molecular switch DnaA-ATP as a central integrator of the bacterial cell cycle.

## RESULTS

### DnaA impacts cell size without changing oriC copy number

A common strategy to study the essential protein DnaA is to use constructs where the native *dnaA* gene is knockout and a *dnaA* copy is engineered to be expressed from inducible promoters. While characterizing such constructs, we noticed that cells expressing *dnaA* from the inducible promoter vanillate are smaller compared to wildtype cells with *dnaA* at its native locus (Figure 1B). The quantification of cell size revealed that cell length decreased ∼20 % (Figure 1C) whereas the width remained the same (Figure S1A). To ensure that this effect was not in response to the inducer vanillate, we analyzed cells expressing *dnaA* from the other commonly used promoter, *P_xylose_* ^35^. Our data revealed that cells expressing *dnaA* from either inducible promoter displayed the same reduction in cell length (Figure S1B,C).

Given that DnaA initiates replication at *oriC* and *oriC* copy number are correlated with cell size ^36^, we examined the possibility that changes in *oriC* copy number caused the change in cell size. In *C. crescentus*, replication initiates only once per cell cycle resulting in cells with one or a maximum of two *oriC* copies ^34^. To quantify *oriC* copy number, we used the centromere-like region *parS* (∼8 kilobases away from *oriC*) as proxy of number of origins of replication ^37,38^. We fluorescently labeled the *parS*-binding protein ParB and tracked the number of CFP-ParB foci. Our data showed that cells expressing *dnaA* from inducible promoters display wildtype numbers of *oriC* copies per cell (Figure 1D), discarding the possibility that changes in *oriC* copy number are responsible for the effect on cell length.

Taking advantage of *C. crescentus* asymmetric cell cycle, we examined whether the DnaA-dependent impact on cell length was specific to the developmental stage of the cell. We wondered about this possibility based on the recent findings that *C. crescentus* shows differential growth rates through their cell cycle ^39,40^. To test this hypothesis, we analyzed cell size of isolated homogeneous population of swarmer cells ^30,31^. Our analysis revealed that swarmer cells display a similar ∼20% reduction in cell length (Figure 1E) demonstrating that the impact on cell size is not specific to the developmental stage of the cell. Collectively, these data revealed that changing *dnaA’s* transcriptional regulation result in a reduction of cell length independent of both *oriC* copy number and cell cycle stage.

### Cell length inversely correlates with DnaA abundance

The levels and activity of DnaA as the replication initiator are regulated at a multitude of levels, including post-transcriptionally and post-translationally ^41,42^. Given that our observed reduction in cell length was potentially linked to *dnaA*’s transcriptional regulation (native vs. inducible promoter), we examined whether the levels of the protein DnaA had changed when expressed from inducible promoters. Using western blot analysis with antibodies specific to DnaA, we found that the small cells display ∼3-fold higher levels of DnaA compared to wildtype grown under the same condition (Figure S1D,E), suggesting increased DnaA levels were causing changes in cell length. We next examined whether the changes in cell length are proportional with the cellular levels of DnaA under conditions that do not impact *oriC* copy number. We titrated the induction levels of *dnaA* expression and confirmed the corresponding increase in DnaA protein levels using western blot analysis (Figure 1F). Notably, as the levels of DnaA increase, we observed a corresponding decrease in cell length (Figure 1G). We were able to reach the highest expression levels of DnaA (without over-initiation of chromosome replication) when cells were grown with the inducer xylose as the sole carbon source (no glucose added to avoid repression of the xylose promoter ^43^). Our analysis showed that the highest DnaA concentration also results in the highest decrease in cell length (Figure 1H). Collectively, these data demonstrate that the observed changes in cell length are dependent on the cellular levels of the protein DnaA. From here on, we will refer to cells expressing *dnaA* from inducible promoters (*xyl* or *van*) that exhibit a 3-fold increase in DnaA levels as 3x-DnaA cells.

### Cell size control pathways and their relevance to 3x-DnaA cells

Building on our observation that increased DnaA levels reduce cell length, we next examined whether this DnaA-dependent effect intersects with known bacterial cell size control pathways. For instance, reducing the total protein content in the cell has been shown to correlate with a reduction in cell size ^44^. In *B. subtilis,* reducing the synthesis of proteins (using subinhibitory levels of the ribosome targeting antibiotic chloramphenicol) results in ∼10% cell length reduction without changes in cell width ^45^. Analogous to *B. subtilis*, our analysis of wildtype *C. crescentus* cells exposed to sub-lethal concentrations of chloramphenicol revealed a reduction in cell length by approximately 10%, while cell width remained unchanged (Figure S2A). Notably, chloramphenicol treatment further exacerbated the size reduction in 3x-DnaA cells compared do untreated conditions (Figure S2B). These results revealed that DnaA’s effect on cell size is independent from the effect that protein content has on cell size regulation. To assess the impact of additional cell size determinants on the reduced length of 3x-DnaA cells, we performed supplementary analyses using genetic approaches, growth media types, and small molecule inhibitors (see Supplemental Information for experimental details). The results of these experiments ruled out contributions from the stringent response (Figure S2C) and nutrient availability (Figure S3), but left open the possibility of envelope biosynthesis (Figure S4), which is addressed in detail later in this manuscript.

### The ability of DnaA to bind both ATP and DNA are required for its impact on cell size

To gain insights into the mechanism of how DnaA influences cell size, we examined the functional contributions of its distinct domains (Figure 2A) by constructing a set of variants featuring domain-specific truncations or targeted amino acid substitutions. Because DnaA is essential and minor modification to any of these domains can impact viability, we analyzed the effects on cell size using chromosomal *dnaA* merodiploid strains. These merodiploid strains express variants of DnaA from an inducible chromosomal promoter (*P_xyl_*) while the native copy of *dnaA* remains intact. Our data revealed that this type of constructs with wildtype *dnaA* expressed from *Pxyl* (*parB::cfp-parB P_xyl_-dnaA*) also display the ∼20% reduction in cell length when DnaA levels are increased ∼3-fold (Figure S5A, B). We confirmed that the expression of DnaA variants from *P_xyl_* have no dominant negative effect on viability using colony forming units (CFU) (Figure 2B). Expression and accumulation of DnaA protein variants compared to empty vector control (EV) were confirmed with western blots (Figure 2C).

**Figure 2.**
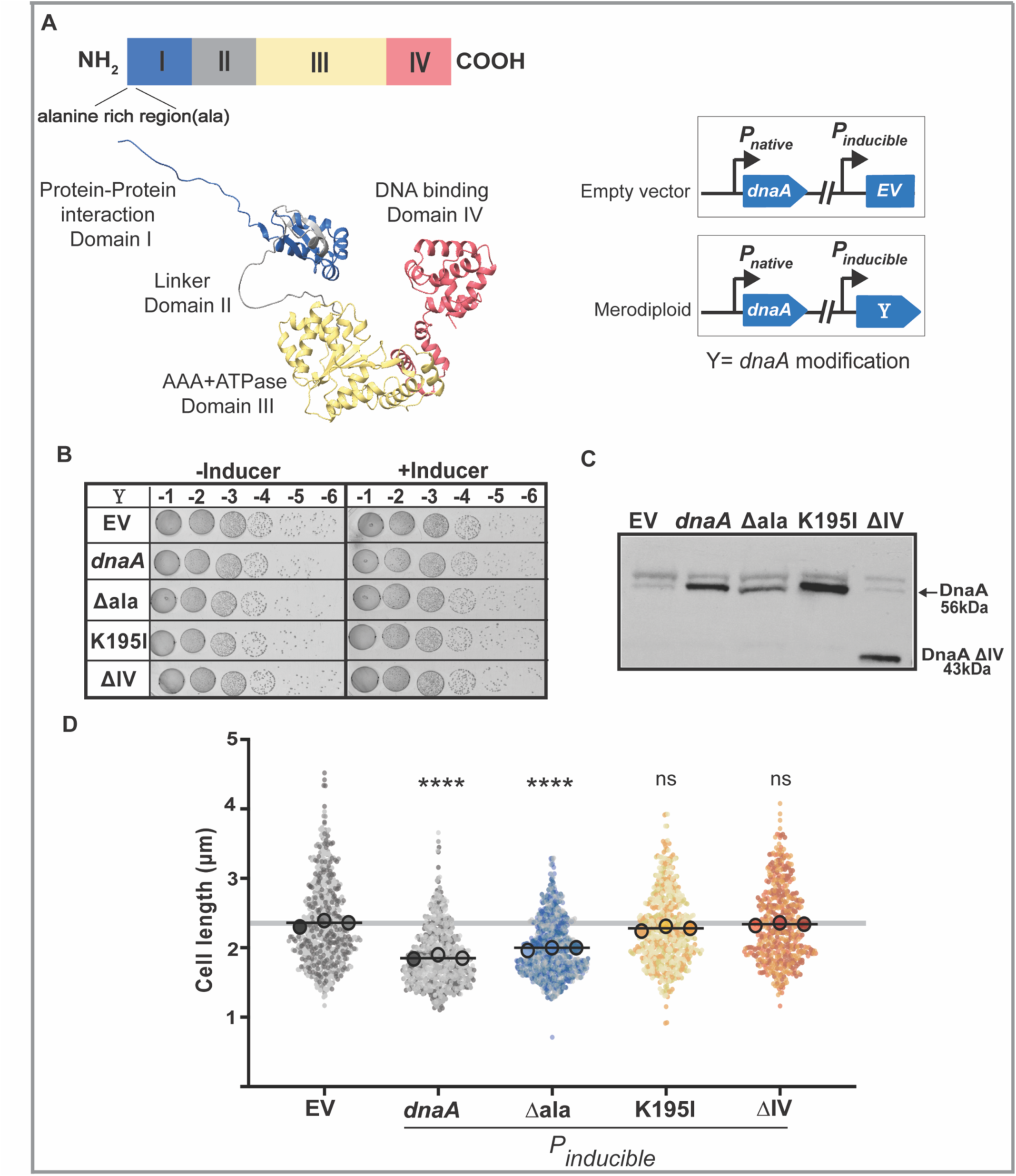
The ability of DnaA to bind ATP and DNA are required for its impact on cell size. **A.** AlphaFold model of DnaA, domains color coded in blue-domain I, grey-domain II, yellow-domain III, red-domain IV. DnaA truncations show WT and mutant DnaA. Schematic shows empty vector (EV) and merodiploid. (merodiploids have a WT *dnaA* under native promoter and *dnaA* modification under *P_xyl_*). Cells were grown overnight in minimal media M2G or M2X (xylose inducer) to exponential phase before serial dilution, western sample preparation and microscopy. **B.** Colony forming units showing viability of DnaA mutants in comparison to empty vector EV (*parB*::CFP-*parB P_xyl_* EV). EV and DnaA mutants show similar growth. **C.** Western blot showing levels of DnaA mutants in comparison to EV. Truncation of domain IV yields 43kDa DnaA. **D.** Super plots showing cell length quantification of EV and DnaA mutants. Small dots represent data points from three independent replicates; large dots represent median values. 3x-DnaA shows ∼20% decrease in cell length. *Δala* rich region from N-terminus (*parB*::CFP-*parB P_xyl_*-Δ*ala* N-ter *dnaA*) shows ∼15% decrease in cell length. DnaA ATP*K195I (*parB*::CFP-*parB P_xyl_-dnaA ^K195I^)* and DnaA domain IV truncation (*parB*::CFP-*parB P_xyl_-dnaA ^I,II,III^*) shows WT cell length ∼2.36µm. Data points show mean ± SD. A parametric t test was performed using population mean values (N = 3). p(<0.0001) ****, p(0.0001)***, p(<0.05)*, ns (non-significant). All samples were blinded n=∼600 cells.

In *C. crescentus*, an alanine-rich region at the N-terminal end of domain I has been shown to regulate the cellular levels of DnaA upon nutrient starvation ^46^. We reasoned that if this ala-rich domain was involved in triggering the change in cell size, increasing the levels of a truncated version of DnaA missing this alanine rich region would have no effect on cell length. However, cells expressing a truncation of the alanine rich region expressed from *P_xyl_* (referred as Δala) retained their ability to reduce their cell length (Figure 2D), albeit to marginally lower levels. The small difference in cell length reduction (∼18% vs. ∼22%) can be attributed to lower levels of expression of this variant compared to cells with 3x-wildtype-DnaA (Figure 2C). These data further support that the impact on cell length is dependent on DnaA levels, and that the alanine rich N-terminal domain is not responsible in DnaA’s regulation of cell length.

We considered another possibility where DnaA’s role with cell size regulation was independent of its ability to bind DNA. We posit that DnaA itself could regulate cell size potentially by directly modulating the activity of cell size regulators independent of DNA binding. We tested this hypothesis by analyzing the effect on cell length from a DnaA with the DNA-binding domain (domain IV) truncated (referred as ΔIV). Expression of this truncated variant unable to bind DNA had no effect on cell length compared to wildtype levels of DnaA (Figure 2D), revealing that DnaA’s ability to bind DNA is required for its role with cell size regulation. Wondering whether ATP binding and/or hydrolysis was required for DnaA’s impact on cell length, we analyzed a variant with a single amino acid substitution that interrupts ATP binding. We did not analyze DnaA variants unable to hydrolyze ATP because such variants are stuck in the active form and over-initiate chromosome replication, which would inevitably impact cell size in an *oriC* copy dependent manner. We constructed DnaA-K195I, a variant encoding a mutation in the Walker A box that affects DnaA’s ability to bind ATP ^47^. Our data revealed that DnaA-K195I lost the effect on cell length and cells expressing this variant displayed wildtype cell size (Figure 2D).

Collectively, the analyses of DnaA variants revealed that DnaA requires to bind DNA and to bind ATP to cause the effect on cell size. Given that DNA-binding and ATP-binding are involved in regulating both functions of DnaA (chromosome replication initiator and transcription factor), we set out to differentiate DnaA’s effect on cell length between these two functions. Because we cannot uncouple these two functions without altering the proper progression of the cell cycle and thus cell growth regulation, we proceeded by analyzing each function independently.

### Chromosome replication remains unchanged in 3x-DnaA cells

The initiation of chromosome replication requires DnaA-ATP oligomerization at *oriC*, which opens the double stranded DNA allowing for components of the replisome to assemble and to initiate replication bidirectionally ^48^. We explored the possibility that 3x-DnaA levels could alter the overall process of chromosome replication, potentially leading to the reduction in cell size. To test this hypothesis, we used time-lapse microscopy to image synchronized swarmer cells throughout their cell cycle, focusing on three key events of chromosome replication: (a) timing of replisome assembly, (b) timing of replication initiation, and (c) progression of the replication forks (Figure 3). To track replisome assembly, we imaged a strain encoding the replisome component DnaN (β-clamp) fluorescently tagged at *dnaN’s* native chromosomal locus ^49,50^. Our analysis of the time when the DnaN-mCherry focus emerges revealed that the 3x-DnaA small cells and wildtype cells assembled their replisomes at similar times (Figure 3A). To determine timing of replication initiation, we tracked the appearance of two *parS* loci (8kb from *oriC*) using the *parS*-binding protein CFP-ParB ^37^. Our data revealed the 3x-DnaA small cells initiate chromosome replication at similar times as wildtype cells (Figure 3B). Lastly, we examined replisome progression by comparing when a chromosomal locus near the gene *pleC* (1.3 Mb from *oriC*) is replicated. To track replication of *pleC*, we used a strain engineered to encode the Yersinia pestis sequence of *parS(*pMT1) near the *pleC* locus and the corresponding ParB(pMT1) fluorescently labelled expressed from an inducible promoter ^51^. We included to this strain a fluorescent tag with the *C. crescentus’* native ParB as an internal control. Analyzing the timing of replication of *parS* and *pleC* revealed no difference between wildtype and 3x-DnaA small cells (Figure 3C). Overall, these data indicate that the higher levels of DnaA in 3x-DnaA small cells do not alter the overall process of chromosome replication.

**Figure 3.**
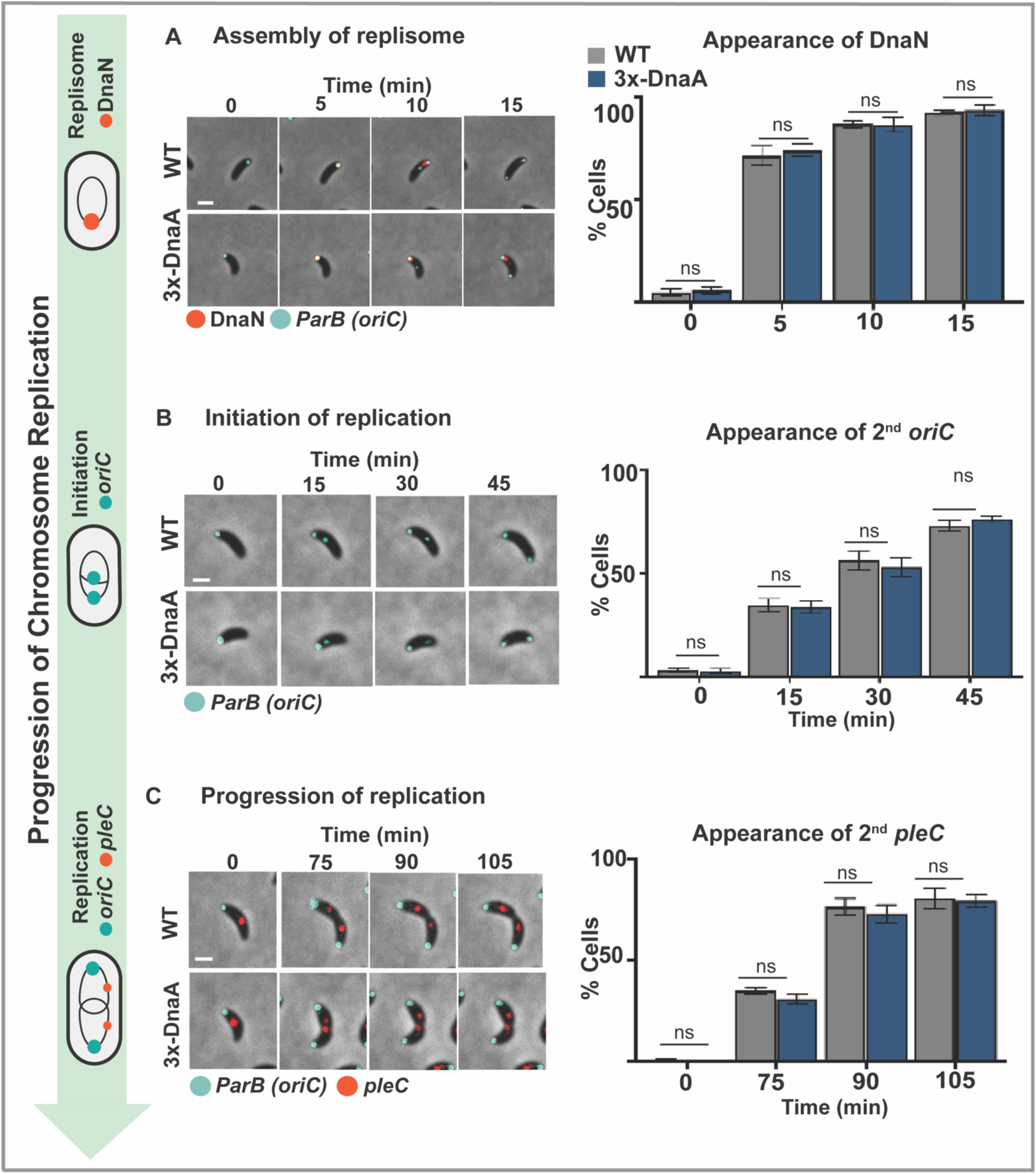
Chromosome replication remains unchanged in 3x-DnaA cells. Cells were grown in minimal media M2G or M2X (xylose inducer) with 100µM van overnight to exponential phase before synchrony. Timelapse images were taken after very 5 minutes for DnaN, every 15min for *oriC* and *pleC*. **A.** Assembly of replisome using fluorescently tagged DnaN in WT (*parB*::CFP-*parB dnaN*::*dnaN* mcherry) and 3x-DnaA (*dnaA*::Ω, *parB::cfp-parB, P_van_ dnaA, dnaN*::*dnaN* mcherry). In both strains replisome assembles at ∼5min represented with red dot. The image is overlayed with CFP *parB* representing *oriC* **B.** Timing of replication initiation was quantified using labelled ParB in WT (*parB*::CFP-*parB*) and 3x-DnaA (*dnaA*::Ω, *parB::cfp-parB, P_van_ dnaA*). In both strains, replication initiation occurs at ∼15-20min, represented with green dot. **C.** Progression of replication using *parS*(pMT1) near the *pleC* locus in WT (CFP-*parB PvanA*-mCherry-*parB*(pMT1) *parS*(pMT1)-*pleC* and 3x-DnaA (CFP-*parB PvanA*-mCherry-*parB*(pMT1) *parS*(pMT1)-*Pxyl-dnaA*). In both strains a second *pleC* tag appears around ∼75min, represented as red dot. Data is representation of three independent experiments. Data points show mean ± SD. A parametric t-test was performed using population mean. p(<0.0001) ****, p(0.0001)***, p(<0.05)*, ns (non-significant). All samples were blinded n=∼100 cells.

### 3x-DnaA cells display changes in transcriptional profiles

After demonstrating that 3x-DnaA levels do not affect the progression of chromosome replication, we proceeded to investigate DnaA’s other conserved role as a master transcriptional regulator. ^52–54^. DnaA transcriptional regulon typically includes genes encoding components of the replisome, cell cycle regulators, and nucleotide biosynthesis ^54^ ^55^ ^56^. To test the hypothesis that DnaA regulates cell size through its activity as a transcription factor, we examined transcriptional changes in 3x-DnaA small cells. For these analyses, we used the *dnaA* merodiploid strains to enable cell growth and comparisons with the DnaA variants. When comparing 3x-DnaA (*P_xyl_-dnaA*^WT^) with an empty-vector control, our RNA-Seq analysis revealed relatively few genes change expression: 45 total genes with 35 upregulated and 10 downregulated (≥1.5-fold change and FDR<0.05). To further exclude non-contributing factors responsible for changes in cell length, we compared RNA-Seq changes between *P_xyl_*-*dnaA*^WT^ cells (exhibit reduced size) and *P_xyl_*-*dnaA-K195I* (exhibit unaffected cell size due to variant’s inability to bind ATP). Using this strategy with the same parameters of ≥1.5-fold change and FDR<0.05, our list of potential genes was reduced to 23 (Figure 4A, Table S1). From this list we focused on genes involved in cell cycle progression. To validate our hits, we used constructs with chromosomal merodiploid strains to circumvent the high expression levels associated with replicating plasmids, given that the genes of interest exhibited changes of approximately 2-fold.

**Figure 4.**
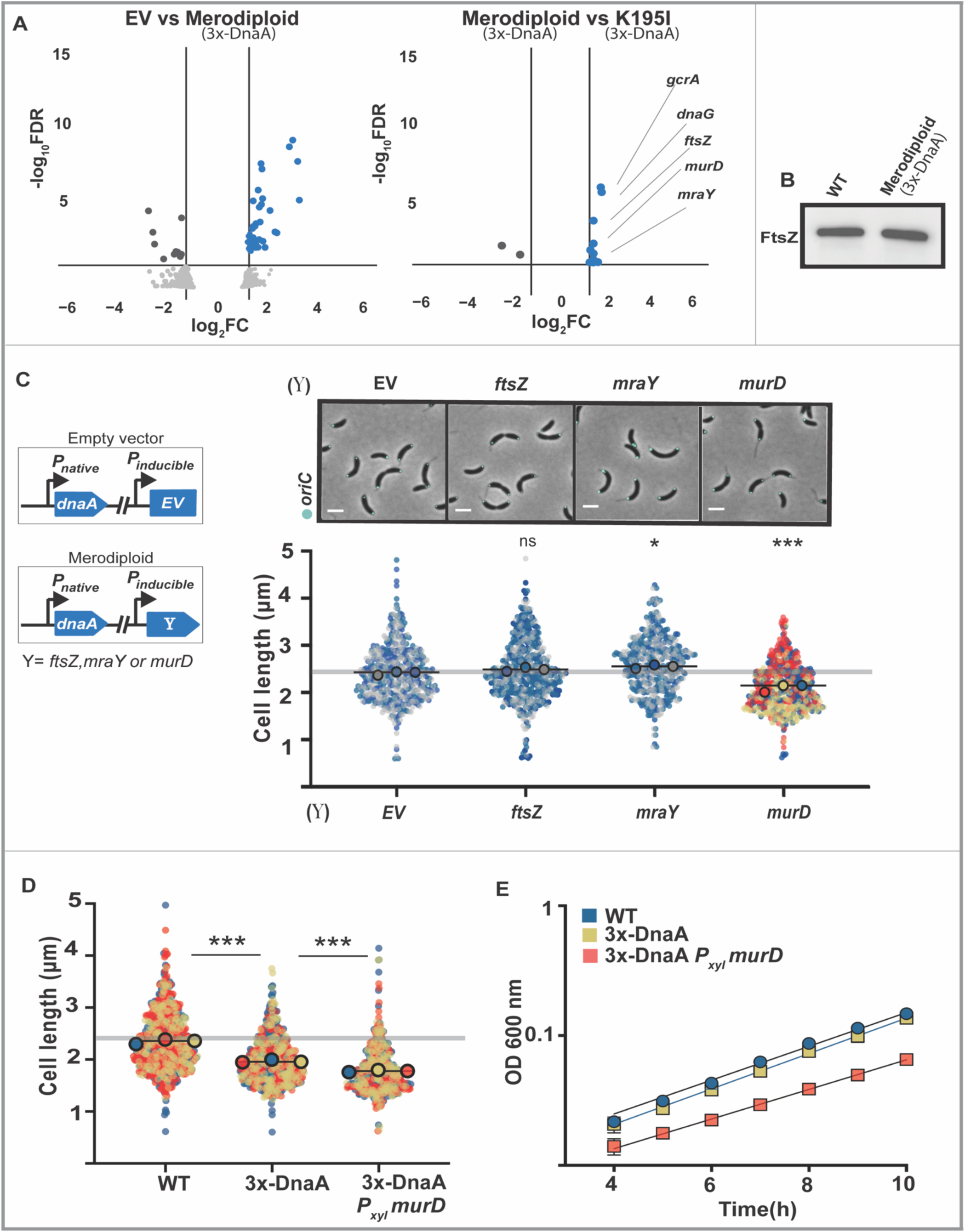
3x-DnaA cells display changes in transcriptional profiles. **A.** Cells were grown in minimal media M2G or M2X (xylose inducer) overnight to exponential phase before RNA extraction, sample preparation and microscopy. **A.** (left) RNA-seq data represented as volcano plot (left panel) comparing EV (*parB*::CFP-*parB P_xyl_* EV) to 3x-DnaA (*parB*::CFP-*parB P_xyl_-dnaA*) represented as dark grey shows 48 genes upregulated and 10 genes downregulated FDR <0.05. (right) Comparison of 3x-DnaA to DnaA K195I (*parB*::CFP-*parB P_xyl_-dnaA ^K195I^*. **B.** FtsZ levels in WT and 3x-DnaA strain shows comparable levels, FtsZ ∼65kDa. **C.** Cell images of EV and merodiploid *ftsZ*, *mraY* and *murD,* scale 2µm. Super plots showing cell length quantification of EV and merodiploid (*parB*::CFP-*parB P_xyl_-ftsZ),* (*parB*::CFP-*parB P_xyl_-mraY),* (*parB*::CFP-*parB P_xyl_-murD).* Small dots represent data points from three independent replicates; large dots represent median values. Induction of *murD* shows 10% decrease in cell length in comparison to EV in minimal media. **D.** Super plots showing cell length quantification of *murD* merodiploid. *murD* overexpression in 3x-DnaA background labelled as 3x-DnaA *P_xyl_-murD* (*dnaA*::Ω, *parB::cfp-parB, P_van_ dnaA, P_xyl_-murD*) results in an additional ∼10% decrease in cell length in comparison to 3x-DnaA (*dnaA*::Ω, *parB::cfp-parB, P_van_ dnaA).***E.** Growth curves showing induction of *murD* in 3x-DnaA cells (3x-DnaA *P_xyl_-murD)* leads to a defect on growth rate. Data is representation of three independent experiments. Data points show mean ± SD. A parametric t-test was performed using population mean. p(<0.0001) ****, p(0.0001)***, p(<0.05)*, ns (non-significant). All samples were blinded n=∼500 cells.

The 1.5-fold increase in *ftsZ* expression in 3x-DnaA cells immediately stood out for multiple reasons. First, *ftsZ* is part of the transcriptional regulon of DnaA ^54^. Second, the transcriptional regulation of *ftsZ* is dependent on the nucleotide-bound to DnaA ^14^. Third, FtsZ is a key regulator of the divisome, the molecular machinery that drives cytokinesis and cell wall biosynthesis at the septum ^57^, both functions that can directly impact cell size ^58,59^. To characterize the potential role of FtsZ, we first examined whether the levels of the FtsZ protein were higher in 3x-DnaA cells compared to empty vector, like our transcriptomics data suggested. However, western blot analysis using FtsZ-specific antibodies revealed no detectible difference in FtsZ protein levels between 3x-DnaA cells and the control (Figure 4B). To further characterize whether elevated levels of FtsZ could lead to a reduction in cell size, we constructed a chromosomal *ftsZ* merodiploid strain that displayed subtle increases in FtsZ levels (Figure S6A). Our data revealed that increasing FtsZ levels does not decrease cell length (Figure 4C). Instead, cells with increased FtsZ levels retain similar size as wildtype but they generate mini-cells (Figure S6B), consistent with previous reports ^60^. These data demonstrate that the DnaA-dependent changes in cell size occur independently of changes in levels of FtsZ.

### MurD is a key regulator that impacts cell size in a DnaA-ATP-dependent manner

The transcriptional analysis of 3x-DnaA cells revealed changes in two peptidoglycan (PG) biosynthetic genes located adjacent to each other: *murD* (+1.5-fold) and *mraY* (+1.6-fold). MurD is one of the four cytoplasmic ATP-dependent enzymes that catalyze the successive additions of amino acids to UDP-N-acetylmuramic acid in PG biosynthesis ^61^. MraY transfers UDP-N-acetylmuramic acid-pentapeptide onto the lipid carrier bactoprenol forming the intermediate known as lipid I ^62^. To test the hypothesis that changes in MurD or MraY levels were responsible for the changes in cell size observed in 3x-DnaA small cells, we constructed chromosomal merodiploid strains of each to titrate the levels of these proteins using wildtype expression of *dnaA* as the parent strain. Induction of *mraY* expression in the *mraY* merodiploid strain did not reduce cell length but instead caused a subtle but significant increase in cell length compared to empty vector (Figure 4C). Remarkably, analysis of the *murD* merodiploid strain revealed that induced increased expression of *murD* alone results in cells with ∼10% reduction in cell length. We show that MurD’s ability to restrict cell length is independent of nutrient availability (Figure S7).

Altering the expression of *murD* was the only genetic modification from our RNA-Seq analysis that led to a decrease in cell size (see other genes tested in Supplemental information and Figures S6) Wondering whether we could achieve an even greater reduction in cell size by further increasing *murD* expression beyond the already higher levels in 3x-DnaA cells, we constructed a *murD* merodiploid strain using the 3x-DnaA cells as the parent strain (*dnaA* expressed from inducible promoter). Our data revealed that indeed, induction of *murD* overexpression in the 3x-DnaA cells results in an additional ∼10% decrease in cell length generating even smaller cells than 3x-DnaA cells (Figure 4D). However, the higher induction of *murD* in 3x-DnaA cells and/or their smaller cell size led to a defect in doubling rate (Figure 4E), demonstrating the importance of maintaining proper levels of MurD and cell size for optimal growth.

### DnaA-ATP regulates the transcription of murD

Having identified MurD as a key factor in regulating size in 3x-DnaA cells, we next examined the relationship between *murD* expression and DnaA. We initially hypothesized that DnaA regulates *murD* expression indirectly through the transcription factor GcrA ^25^. This hypothesis was based on two observations: first, gcrA transcript levels were elevated 1.8-fold in 3x-DnaA cells (Figure 4A); second, *murD* is part of the division and cell wall (dcw) cluster ^18^, and in *C. crescentus,* its expression has been predicted to be under GcrA control ^63^. To test our hypothesis, we modulated GcrA levels independently of DnaA and quantified *murD* transcripts using RT-qPCR. These analyses revealed that changes in GcrA abundance had no detectable effect on *murD* expression (Figure 5A). In contrast, altering DnaA levels result in clear changes in *murD* transcript levels (Figure 5B), suggesting a direct regulatory role for DnaA.

**Figure 5.**
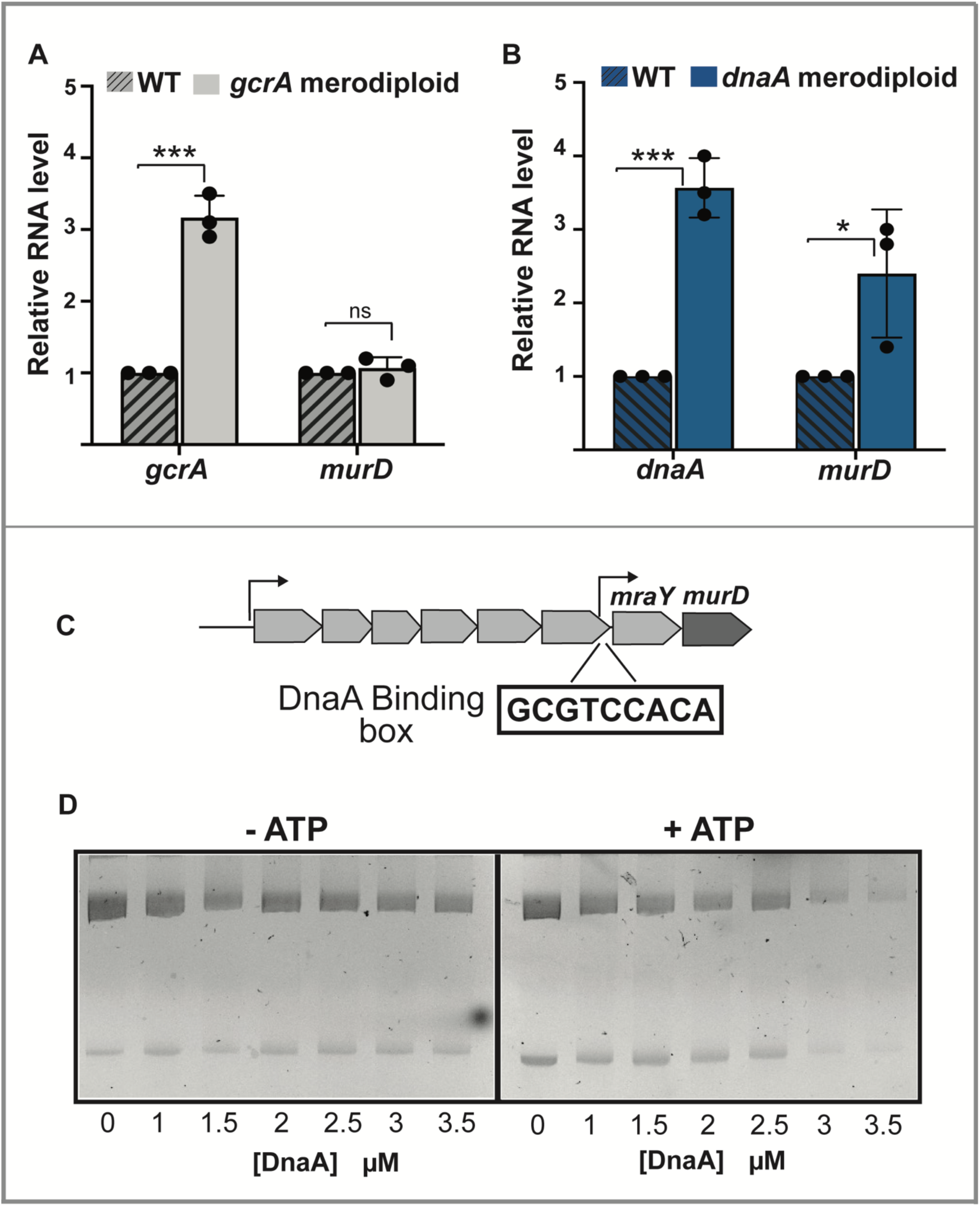
DnaA-ATP regulates the transcription of *murD*. **A.** (left) RNA levels of *gcrA* and *murD* in WT (*parB*::CFP-*parB)* in comparison to *gcrA* merodiploid (*parB*::CFP-*parB P_xyl_-gcrA). gcrA* RNA levels were increased in *gcrA* merodiploid strain in comparison to WT. *murD* levels were non-significantly changed in comparison to WT. **B.** (right) RNA levels of *dnaA* and *murD* in WT (*parB*::CFP-*parB)* in comparison to *dnaA* merodiploid (*parB*::CFP-*parB P_xyl_-dnaA). dnaA* RNA levels were increased in *dnaA* merodiploid strain in comparison to WT. *murD* levels were significantly increased *dnaA* merodiploid strain. p(<0.0001) ****, p(0.0001)***, p(<0.05)*, ns (non-significant). **C.** Schematic of *murD* operon in *Caulobacter*, with another +1 site upstream of *murD.* Strong DnaA binding box was identified upstream of *mraY.* **D.** EMSA showing DnaA binding with purified DnaA protein. 200bp upstream of *mraY* containing the DnaA binding box were cloned in a plasmid and used as DNA. Purified DnaA + DNA was incubated with ATP without ATP 30° for 10 minutes. Increasing DnaA concentration in presence of ATP shows a shift, highlighting DnaA binding to the DnaA binding box in the presence of nucleotide.

Given that *murD* is part of an operon (Figure 5C), we were intrigued that only *mraY* and *murD* were significantly upregulated in 3x-DnaA cells (Figure 4A). This selective expression suggested that these genes are transcribed independently of the rest of the operon. Supporting this hypothesis, previously reported +1 transcriptional start sites (TSS) were identified upstream of the *mraY* gene^64^. Furthermore, we identified a DnaA binding motif upstream of the translational start codon of *mraY* (Figure 5C), with structural organization previously shown with other genes regulated by DnaA ^54^. To directly test the model that DnaA can bind upstream of *mraY-murD*, we cloned the region upstream of *mraY* into a plasmid and tested direct binding using electrophoretic mobility shift assays (EMSA) with purified DnaA^65^.

Our EMSA analysis confirmed that DnaA binds upstream of *mraY-murD*, supporting DnaA’s role as a transcriptional regulator (Figure 5D). Furthermore, we discovered that DnaA’s ability to bind upstream of *mraY-murD* is significantly increased when DnaA is bound to ATP (Figure 5D), indicating that nucleotide binding is essential for DnaA’s regulatory activity of *mraY-murD* expression. This ATP requirement is remarkably consistent with our analyses of cells expressing the ATP-binding deficient variant DnaA-K195I that was unable to influence cell size (Figure 2D) and displayed no differences in *mraY* and *murD* transcriptional levels compared to wildtype or to 3x-DnaA cells (Table S1). These findings reinforce the essential role of ATP in enabling DnaA to function as a transcriptional regulator of cell wall biosynthesis.

### 3x-DnaA small cells constrict at a faster rate during cell division than WT

Constriction rate plays a critical role in the regulation of cell size in bacteria ^28^. Given that our 3x-DnaA cells display smaller size, we investigated whether the reduction in size could be attributed to a change in constriction dynamics. To determine rate of constriction, we imaged synchronized 3x-DnaA cells and wildtype cells through a complete cell cycle starting from the G1 phase (swarmer state) through the completion of cytokinesis. Each individual cell was then analyzed through the progression of its cell cycle using the Fiji plugin MicrobeJ ^66,67^. Using automated 5-min interval time-lapse image data, our analysis revealed that 3x-DnaA cells constrict at a significantly faster rate (10.1 ± 0.6 nm/min) than wildtype cells (7.4 ± 0.5 nm/min), representing an increase of approximately 26% (Figure 6C). In *C. crescentus,* faster constriction rates are correlated with slower elongation rates ^68–70^. Our analysis of elongation rate demonstrated that, 3x-DnaA cells elongate at a slower rate (8.1 ± 0.7 nm/min) compared to wildtype cells (10.7 ± 1.0 nm/min) (Figure 6D). Collectively, these findings support that DnaA’s transcriptional regulation of PG biosynthetic enzymes can modulate cell size by altering the cell growth and division dynamics.

**Figure 6.**
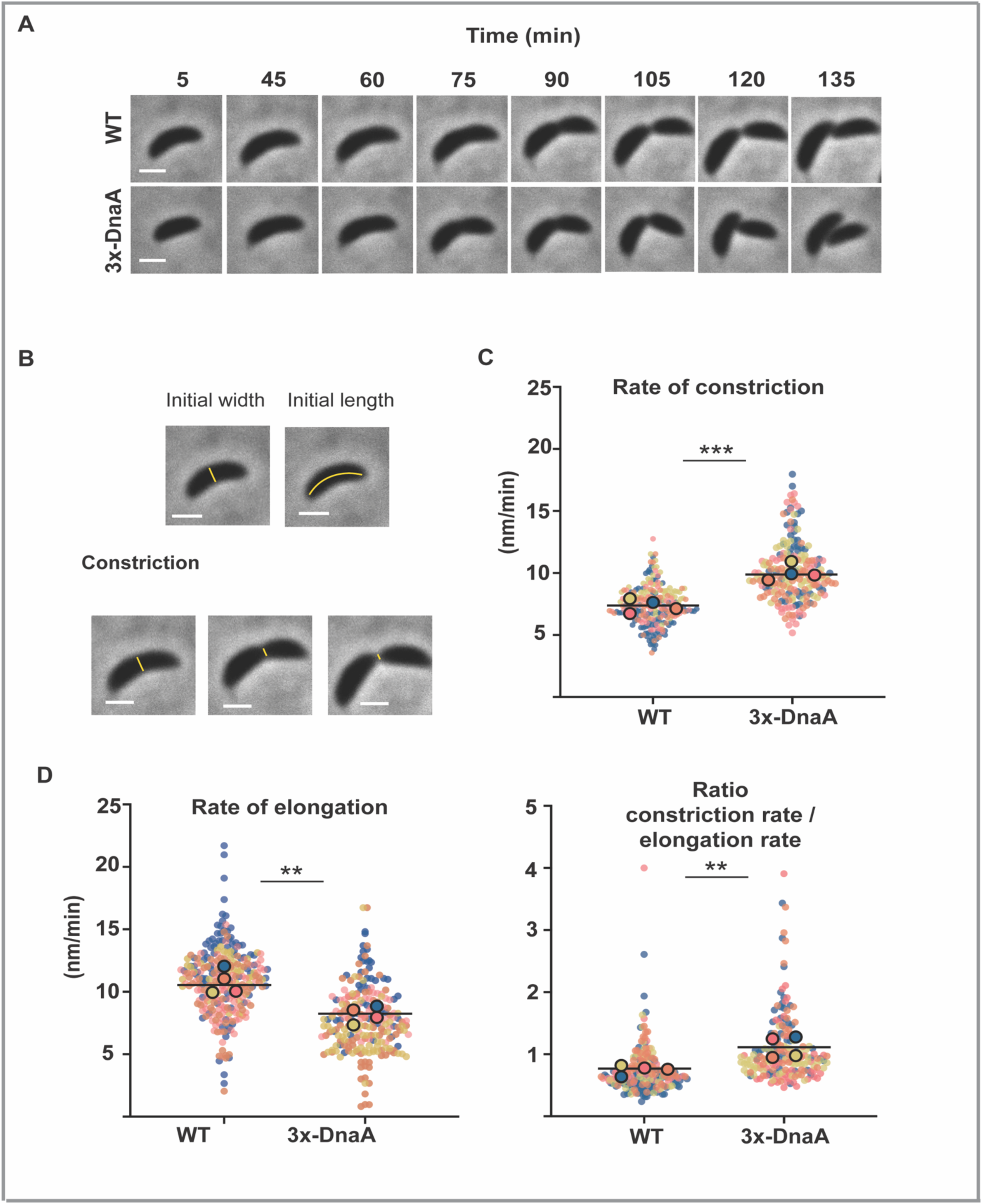
DnaA modulates cell growth and constriction rates. **A.** Timelapse phase contrast images of WT (*parB*::CFP-*parB)* and 3x-DnaA (*dnaA*::Ω, *parB::cfp-parB, P_van_ dnaA)*. Cells were grown in M2G overnight with inducer 100µM van to exponential phase before synchrony, time-lapse images were taken after very 5 minutes, representative images are shown at 5 min, and then 45 min to 135 minutes post synchrony. scale 2µm. **B.** Phase contrast images showing initial length and initial width and constriction. scale 2µm. The yellow lines represent measures of length or width for each cell. **C.** Super plots showing quantification of rate of constriction in WT and 3x-DnaA. Constriction was significantly faster in 3x-DnaA cells. **D.** Super plots of quantification of rate of elongation shows it is reduced by ∼20% in 3x-DnaA cells. Super plot of ratio of rate of constriction over rate of elongation shows a shift in favor of constriction in 3x-DnaA cells. Small dots represent data points from four independent replicates, large dots represent median values (blue,pink,orange,yellow). Each data point is an independent experiment. Data points show mean ± SD. A parametric t-test was performed using population mean values (N = 4) to compare values for each measurement. p(<0.0001) ****, p(0.0001)***, p(<0.05)*, ns (non-significant). All samples were blinded n= ∼300 cells.

## DISCUSSION

In this study, we investigated how elevated levels of DnaA, without disrupting chromosome replication, result in a consistent ∼20% reduction in cell size. Using the genetically tractable model *C. crescentus*, we dissected the overlapping events of the bacterial cell cycle and found that ATP-bound DnaA not only initiates chromosome replication but also influences cell growth and constriction rate by regulating the transcription of genes involved in cell wall biosynthesis. Notably, we identified MurD, a conserved enzyme in cell wall biosynthesis, as a previously unrecognized determinant that restricts cell size. Our data support the model that DnaA functions as a transcriptional regulator of *murD*, and that this regulatory activity is dependent on its ATP-bound state.

We propose that DnaA orchestrates a broad transcriptional program, integrating modest shifts across multiple targets to coordinate cell growth. While increased *murD* expression alone led to ∼10% reduction in cell size, increased DnaA levels produced a more pronounced ∼20% decrease. Our transcriptional analyses revealed subtle (<2-fold) changes in genes involved in cell wall biosynthesis and cell cycle progression in cells with increased DnaA levels (Table S1). Although none of these genes individually recapitulated the full phenotype and most caused cell filamentation when perturbed (Figure S6), their collective regulation likely contributes to the observed size reduction. This is consistent with the complexity of cell growth, which requires the synchronized activity of diverse enzymes and protein complexes.

Bacteria have been known to follow the Growth law, which states that cell size is an exponential function of growth rate ^71–75^. In this study, we show that MurD’s impact on cell size is independent of nutrient availability (Figure S7). Cells with increased *murD* expression in wildtype background reduced their size when grown in minimal or rich media. We discovered that further increase of *murD* expression results in a further reduction in cell size that was interestingly accompanied by a decrease in growth rate. This phenomenon may be attributed to the reduced cellular volume interfering with essential housekeeping processes during the cell cycle or a potential connection between MurD and the Growth Law. MurD has gathered significant attention in drug development due to its crucial role in cell wall biosynthesis ^76,77^. Further investigation into MurD’s activity and its role in cell size regulation will yield significant insights into the complex mechanisms that govern the progression of the bacterial cell cycle.

The link between rates of growth/constriction and enzymes involved in the biosynthesis of PG precursors has been observed across different bacterial species^78^. In *B. subtilis*, the growth rate is dependent on the abundance of lipid II and of the levels of the first committed PG biosynthetic enzyme, MurA ^79^. In *P. aeruginosa*, MraY, the enzyme that synthesizes Lipid I, maintains a balance in cell wall homeostasis and growth ^80^. In *Staphylococcus aureus*, MurJ, the membrane-bound protein that flips Lipid II from the cytoplasm to the periplasm, has been linked to accelerating constriction rate independent of FtsZ-treadmilling ^81^ ^82^. In *C. crescentus*, a positive correlation was shown between constriction rate and PG proteins FtsWI ^68^ and their regulator FzlA ^70 83^. The work presented here demonstrates that the PG biosynthetic enzyme MurD is also linked to the regulation of growth/constriction rates. Furthermore, our data uncovered an additional layer of regulation that integrates chromosome replication into the coordination between PG biosynthesis and growth/division, underscoring the sophisticated mechanisms bacteria employ to sustain rapid and accurate proliferation.

The discovery that DnaA modulates cell growth and division adds to its growing reputation as a global regulator of bacterial cell development, extending beyond its role with chromosome replication. For instance, DnaA’s transcriptional regulon commonly includes other global regulators involved in genome maintenance, cell development, and pathogenesis ^54,84,85^. In *B. subtilis*, DnaA is connected to the onset of sporulation by transcriptionally regulating an inhibitor of the sporulation master transcription factor SpoOA^53^. In *Borrelia burgdorferi*, DnaA regulates the transcription of the nucleoid associated protein EbfC which itself regulates the transcription of a multitude of other genes ^84,86^. Furthermore, altering the transcriptional activity of DnaA in *B. burgdorferi* led to changes in >20 components of the elongasome and divisome, which could also connect DnaA to cell wall biosynthesis ^87^. Beyond bacteria, replication initiators in eukaryotes (Origin Recognition Complex, ORC) also perform roles beyond DNA replication, influencing multiple cellular events including chromosome repair, condensation, segregation, and cytokinesis^88–92^. These functional parallels across domains of life emphasize the role that chromosome replication initiators play as central integrators of the cell cycle.

## RESOURCE AVAILABILITY

All data are contained within the manuscript.

## ACKNOWLEDGEMENTS

We are grateful to Prof. Erin Goley and her lab for providing the FtsZ antibodies, specifically to Dr. Jordan M. Barrows for his assistance with the rate of elongation and constriction protocol. We thank Anuradha Sharma from the Prof. Sanfilippo lab for her help with the RT-qPCR, Dr. Alvaro Hernandez from the Roy J. Carver Biotechnology Center for RNA sequencing, as well as to Dr. Jenny Drnevich for her expertise in the statistical analysis of the RNA-seq data. We thank Sasha Agafonova for her contribution in the construction of the graphical abstract. The work reported in this publication was supported by the National Institute of General Medical Sciences of the National Institutes of Health (NIH) under Award Numbers R01 GM133833 & R3GM158016, and by the National Science Foundation (NSF) Science and Technology Center for Quantitative Cell Biology under Award Number 2243257.

## AUTHOR CONTRIBUTIONS

EK and PEM conceived project and wrote manuscript. EK performed all experiments and data analyses.

## MATERIALS AND METHODS

### Bacterial strains and growth conditions

Strain and plasmid descriptions are listed in the Supplementary Table S2, S3. *Caulobacter crescentus* strains used in this study were derived from NA1000 (wildtype). Plasmids were constructed by cloning PCR products into pNPTS138,pXCHYC-2, pBXMCS-2 vectors ^35^. Amplified PCR products were placed in vectors via Gibson or restriction cloning. Plasmids were transformed into DH5α cells and verified through sequencing. In *C. crescentus*, plasmids were transformed via electroporation. *Caulobacter* strains were grown in peptone yeast extract (PYE) or M2 minimal media supplemented with 0.2% glucose (M2G) or 0.2% xylose (M2X). Liquid cultures were re-inoculated overnight in fresh media to grow cells until exponential phase OD_600_ ∼ 0.3-0.4 at 30°C under mechanical agitation 200 rpm. Overnight cultures were supplemented with specific antibiotics and inducers. Vanillate 100µM was used to induce the expression of *P_van_*, while 0.3% xylose or M2X was used to induce *P_xyl_*. Liquid cultures were supplemented with following concentrations of antibiotics: 5µg/ml kanamycin, 25µg/ml spectinomycin, 5µg/ml streptomycin. For PYE plates 25µg/ml kanamycin, 100µg/ml spectinomycin and 5µg/ml streptomycin. To check the impact of chloramphenicol, cerulenin and fosfomycin, 1/10 (2 µg/ml) sub-lethal concentration of chloramphenicol, 0.6µg/ml cerulenin and 5µg/ml fosfomycin was added to liquid culture and left overnight such that the OD is 0.3 next day.

### Synchronization

*C.crescentus* cells were synchronized using mini-synchrony protocol to isolate swarmer population ^93^. Cells were inoculated in 15ml M2G overnight, to an OD_600_ ∼ 0.3. Cultures were supplemented with inducer and antibiotics as noted. Cells were pelleted at 6000 rpm for 10 minutes at 4°C in the centrifuge. Cell pellet was resuspended in 900µl of 1X M2 salts and 900µl of Percoll (Sigma-Aldrich) for density gradient. The mixed solution was centrifuged at 11.000rpm for 20minutes, the bottom layer of swarmer cells was isolated and transferred to a fresh tube. Cells were washed twice with ice cold 1X M2 salts, centrifuged at 8000 rpm for 3 minutes at 4°C. Swarmer’s were resuspended in M2G to appropriate OD_600_ ∼0.2-0.3.

### Growth assays

*C.cresentus* overnight cultures were grown from freezer stock in M2G or PYE liquid medium were back diluted to reach OD_600_ ∼0.1 for growth curves. Early log phase cultures were used for inoculation in 96-well plate in M2G or PYE with relevant inducer. Optical density at 600 nm was monitored every hour at 30°C in a Biotek EPOCH-2 microplate reader with shaking.

### Microscopy

Cells were grown to an exponential OD_600_ ∼0.3. Cells (1µl) were spotted on agar pads (1% agarose in M2G or M2salts). For time-lapses (2µl) cells were spotted on agar pads. Phase contrast and fluorescent images were taken at room temperature using Zeiss Axio Observer 2.1 inverted microscope with AxioCam 506 mono camera (objective: Plan-apochromat 100x/1.40 Oil Ph3 M27 [WD=0.17mm]) and Zen Pro software. Samples were blinded for data analysis. To count number of foci, ImageJ (cell counter plugin) was used ^94^. Cell size (length and width) was analyzed using ImageJ/FIJI plugin MicrobeJ ^67^. Multiple frames were analyzed for cell size analysis, data were presented in Superplots ^95^.

### Elongation and constriction

Cells were grown in liquid cultures with inducer overnight to reach early log phase OD_600_ ∼0.3 to synchronize. After synchrony cell were resuspended in M2G or M2X with appropriate additives. Cells were then spotted on agar pads (0.1% agar with inducer) for timelapse. Images were taken after every 5 minutes through one cell cycle. MicrobeJ was used to analyze timelapse images as described by ^83,96^. Constriction is automatically detected at the positive curvature midcell and manually segmented upon division. Cell width was detected at the site of constriction whereas cell length was calculated at each time point. Cell width was detected at the site of constriction whereas cell length was calculated at each time point. Constriction time was determined by multiplying the number of frames from constriction initiation to division by 5 (images were taken every 5min). The constriction rate was calculated by dividing the cell width at the onset of constriction by the constriction time. Elongation rate calculated by dividing change in length from constriction initiation to division by constriction time.

### Immunoblotting

*Caulobacter* mixed population cells were grown to early exponential phase OD_600_ ∼0.3 with inducer. The OD_600_ of the incubated cultures was normalized to 0.2. Cells were pelted and resuspended in 40µl of cracking buffer, then boiled for 10 min at 90°C. Samples were stored at -20°C for western blotting. A 12% SDS-PAGE gel was used to separate proteins and then transferred to PVD membrane using iBlot2 (Invitrogen Dry Blotting System). The membrane was blocked with 1XTBS (10mM Tris-HCL, pH8,150mM NaCl, 0.1% Tween-20), 0.1% tween 20 (TBST) and 5% non-fat milk at room temperature. The blot was incubated overnight with primary antibody for DnaA (α-DnaA) ^97^ at a dilution of 1:15,000 for FtsZ (α-FtsZ)^98^ at a 1:20,000 dilution in TBST with 5% milk overnight at 4°C. Next day the membrane was washed 3X with TBST and then incubated with secondary antibody (α-Rabbit IgG peroxidase, Sigma-Aldrich) diluted to 1:15,000 in TBST with 5% milk at room temperature for 1 hour. The blot was washed 3X with TBST for 5 minutes. To develop the blot, SuperSignal West Pico PLUS Chemiluminescent Substrate (Themo Fisher Scientific) was used and imaged by using ChemiDoc-MP (Bio-Rad). WT DnaA runs at 56kDa, whereas WT FtsZ is 54.1kDa, but it runs at ∼65kDa.

### CFUs for viability assay

*Caulobacter* cells were grown from freezer stocks in minimal media M2G or M2X, it was then reinoculated in fresh media with relevant additives at 30°C 180rpm to reach exponential growth OD_600_ ∼0.3. Cells were then normalized to OD_600_ ∼0.1. Cultures were serially diluted in a 96-well plate and plated on M2G or M2X plates (0.2% xylose in M2X plates was used as inducer). Plates were incubated at 30°C for 2 days and then imaged using ChemiDoc-MP (Bio-Rad).

### RNA-sequencing

*Caulobacter* cells were grown from freezer stocks in minimal media M2G, reinoculated in fresh media M2G or M2X (M2X media to induce the expression of mutants) at 30°C 180rpm to reach exponential growth OD_600_ ∼0.3. Total RNA was extracted using hot phenol procedure ^99^. After total RNA extraction it was treated with DNase (Invitrogen TURBO DNase) to remove genomic DNA and quantitated using nanodrop (Themo Fisher Scientific NanoDrop One^c^). Samples were then sent to Roy J. Carver Biotechnology Center Sequencing core at UIUC to check RNA integrity through Bioanalyzer, library preparation, sequencing, and data analysis.

### RT-qPCR

*Caulobacter* cells were grown from freezer stocks in minimal media M2G, reinoculated in fresh media M2G or M2X (M2X media to induce the expression of mutants) at 30°C 180rpm to reach exponential growth OD_600_ ∼0.3. Total RNA was extracted using hot phenol procedure ^99^. After total RNA extraction it was treated with DNase (Invitrogen TURBO DNase) to remove genomic DNA and quantitated using nanodrop (Themo Fisher Scientific NanoDrop One^c^). cDNA synthesis and Rt-qPCR was performed using Luna universal one-step RT-qPCR kit. Final concentration of RNA was 10ng for each reaction. To quantify transcription *ruvA* (*CCNA_03345*) was used as an endogenous control, with the following primers: 5′-cgagtgaggaagccgtagag-3′ and 5′-gaccctgttgcacatcgag-3′ ^100^.

### EMSA

EMSA experiments were performed as described previously. DNA fragment was incubated with purified DnaA protein in EMSA buffer containing 20mM HEPES,10mM MgAc, 60mM NaCl, 5% glycerol, 0.1mg/ml bovine serum albumin and 3mM ATP or without ATP. DNA fragment and purified DnaA was incubated in EMSA buffer at 30° for 10 minutes. Samples were run on 2% TBE gel in TBE buffer at 90 V for 6 hours in 4°. DNA was visualized by ethidium bromide staining.

