## Supplemental Data for "The bacterial molecular switch DnaA-ATP integrates chromosome replication with cell growth and division dynamics"

### *DnaA impacts cell size without changing oriC copy number*

Cells expressing *dnaA* from the inducible promoter vanillate are smaller compared to wildtype cells with *dnaA* at its native locus. Quantification of cell size revealed that cell length decreased ~20 % whereas the width remained the same (Figure S1A). To ensure that this phenotype is not promoter specific we analyzed cells expressing *dnaA* from the other commonly used promoter, $P_{xylose}$ [1]. Our data revealed that cells expressing *dnaA* from either inducible promoter displayed the same reduction in cell length (Figure S1B).

### *Cell size control pathways and their relevance to 3x-DnaA cells.*

*ppGpp and cell size* The alarmone molecule guanosine tetra- or pentaphosphate (p)ppGpp, a global inhibitor of biosynthesis, has been implicated in cell size in various bacterial species. The cellular levels of (p)ppGpp are regulated by the Rsh family enzymes RelA & SpoT [2,3]. Unlike *E. coli* and *B. subtilis*, *C. crescentus* and other alphaproteobacterial species only encode the SpoT-dependent pathway [4] In *E. coli*, levels of (p)ppGpp have been shown to impact cell size upon amino acid starvation [5]. In *C. crescentus*, artificially increasing the levels of (p)ppGpp by expressing a constitutively active RelA from *E. coli* results in cells with decreased cell size [6]. To examine whether DnaA's impact on cell length was connected to levels of (p)ppGpp, we constructed a mutant strain unable to synthesize (p)ppGpp by knocking out the gene encoding the bifunctional (p)ppGpp synthetase/hydrolase SpoT. Our analysis revealed that wildtype cells with and without *spoT* retained the same cell size (Figure S2C). Similar to wildtype cells, 3x-DnaA cells with or without *spoT* (*dnaA*::Ω, *parB::cfp-parB*, $P_{van}$-*dnaA*, *ΔspoT)* display the same cell size demonstrating that the alarmone molecule (p)ppGpp is not connected to the observed DnaA-dependent change in size.

*Nutrient availability and cell size* In *C. crescentus*, the DnaA levels are 3- to 4-fold higher when grown in rich media than in minimal media [7-9] . Western quantification in cells expressing *dnaA* from inducible promoters grown in rich media revealed that DnaA levels remained the same as wildtype cells grown in rich media (Figure S3A). Consistent with levels of DnaA impacting cell size, cells expressing *dnaA* from inducible promoters grown in rich media retained the same cell length as wildtype cells grown in rich media (Figure S3B). To increase the levels of DnaA, we constructed two chromosomal *dnaA* merodiploid strains: one with *dnaA* under its native locus plus a copy under an inducible promoter ($P_{van}$-*dnaA*) or a strain with two copies of *dnaA* under two different inducible promoters ($P_{van}$-*dnaA* $P_{xyl}$-*dnaA*). The quantification of DnaA levels with western blots revealed that none of the chromosomal *dnaA* merodiploid strains grown in rich media produced higher levels of DnaA than wildtype. To achieve higher levels of *dnaA* overexpression, we constructed a *C. crescentus* strain with *dnaA* overexpressed from a replicating plasmid [10]. Considering that this approach can lead to over-initiation of chromosome replication, we induced expression of *dnaA* for only ~2 cell cycles in rich media (3h induction). Using western blots, we confirmed that 3h induction results in ~3-fold higher DnaA levels compared to wildtype (Figure S3C). However, even under this short-term induction, we observed a significant percent of cells (>10%) over-initiated chromosome replication (Figure S3D, E). To avoid the implications that over-initiation of chromosome replication has on cell size, we continued our analysis using only chromosomal inducible promoters and only minimal media as the growth condition.

*Envelope biosynthesis and cell size* The biosynthesis of fatty acid and cell wall have also been implicated with cell size regulation. To examine whether fatty acid and peptidoglycan (PG) biosynthesis were linked to the reduction in size of inducible *dnaA (*3x-DnaA) cells, we followed established protocols to obstruct their biosynthesis using sub-lethal concentrations of the antibiotics cerulenin and Fosfomycin. Cerulenin inhibits the first condensation reaction between acetyl-CoA and malonyl-acyl-carrier-protein catalyzed by FabH (b-ketoacyl-acyl-carrier protein synthase) [11]. In *B. subtilis* and *E. coli*, exposure to cerulenin result in ~10% reduction in cell length [12,13]. Unfortunately, we are unable to make any conclusions about cerulenin because we found that the viability of *C. crescentus* is highly sensitive to even low levels of cerulenin (Figure S4A). One potential explanation to this high sensitivity is that *C. crescentus* encodes a single copy of FabH whereas *B. subtilis* has two FabH homologs FabHA and FabHB [14,15]. Regarding PG biosynthesis, Fosfomycin is used to inhibit the first committed enzyme, MurA (UDP-N-acetylglucosamine enolpyruvyl transferase) [16]. Fosfomycin exposure in wildtype *C. crescentus* cells has been reported to cause cells to increase length and width [17,18]. When we analyzed wildtype cells exposed to similar levels of Fosfomycin (5 μg/mL), we found that indeed cells become bigger in length and width (Figure S4B). However, our analysis of *oriC* copy number in both wildtype and 3x-DnaA cells exposed to Fosfomycin revealed that these larger cells over-initiate chromosome replication resulting in abnormal >2 *oriC* copies per cell (Figure S4C). Because we cannot resolve whether multiple *oriC* copies caused the cell size increase or cell size increase caused the over-initiation of chromosome replication, we cannot make any conclusions based on these data about the potential connection between DnaA and PG biosynthesis.

*The ability of DnaA to bind ATP and DNA are required for its impact on cell size* DnaA is essential and minor modification to any of its domains can impact viability, we analyzed the effects on cell size using chromosomal *dnaA* merodiploid strains. These merodiploid strains express variants of DnaA from an inducible chromosomal promoter ($P_{xyl}$) while the native copy of *dnaA* remains intact. Our data revealed that this type of constructs with wildtype *dnaA* expressed from *Pxyl* (*parB::cfp-parB* $P_{xyl}$-*dnaA*) labelled as (3x-DnaA) also display the ~20% reduction in cell length when DnaA levels are increased ~3-fold (Figure S5A,B).

***3x-DnaA cells display changes in transcriptional profiles***

*gcrA*

We next explored an alternative hypothesis where DnaA regulates cell size indirectly through the transcriptional regulon of GcrA. To test whether increased levels of GcrA cause cells to reduce their cell length, we constructed a *gcrA* chromosomal merodiploid strain (*parB::cfp-parB,* $P_{xyl}$-*gcrA*). Increased expression of the master transcription factor GcrA resulted in cells with longer cell length, instead of shorter (Figure S6C), consistent with previous reports [19,20].

*dnaG*

The DNA primase DnaG represented an interesting target to regulate cell size because DnaG directly regulates the rate of DNA replication in a ppGpp-dependent manner [21,22]. To test DnaG's potential role with cell size regulation, we analyzed a *dnaG* chromosomal merodiploid strain (*parB::cfp-parB,* $P_{xyl}$-*dnaG*) and found that induction of *dnaG* results in no changes in cell size (Figure S6C). These results are consistent with our findings that ppGpp and the overall process of DNA replication are not involved in the cell size reduction of 3x-DnaA cells.

*relB-4*

We also investigated the genes that were downregulated in RNA-seq data. Anti-toxin protein *relB-4* was 2.9-fold downregulated in 3x-DnaA cells. To test whether decreased levels of *relB-4* cause cells to reduce their cell length, we analyzed a *relB-4* chromosomal merodiploid strain in inducible *dnaA* (3x-DnaA) background (*dnaA*::Ω, *parB::cfp-parB,* $P_{van}$ *dnaA)*$P_{xyl}$-*relb-4*). We sought to test whether elevating *relB-4* levels could revert 3x-DnaA cells to WT. We found there was not significant difference in cell length of 3x-DnaA cells when *relB-4* was overexpressed, suggesting *relB-4* is not involved in cell size reduction (Figure S6D)

### ***MurD impact cell size in rich media***

Having identified MurD as a key player in the regulation of cell size, we analyzed whether increasing *murD* expression alone results in cell size changes when cells are grown in rich media. Indeed, our data revealed that induction of *murD* in the merodiploid strain with native levels of DnaA grown in rich media displayed a ~10% reduction in cell length compared to empty vector control without impacting growth rate. (Figure S7C,D). These findings support MurD's involvement in cell size regulation that is independent of growth media type.

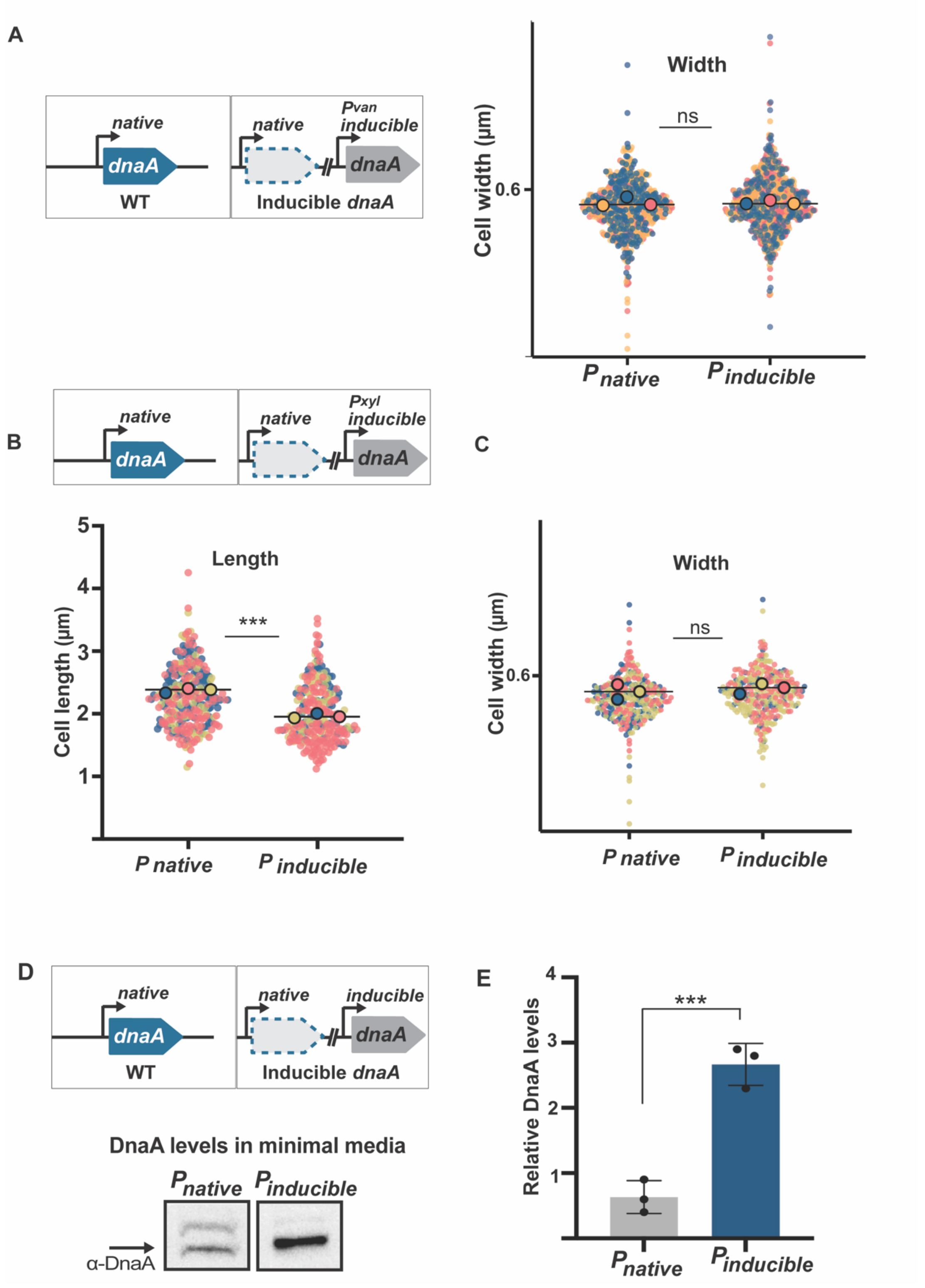


**Figure S1. DnaA impacts cell size without changing *oriC* copy number. A.** WT ($P_{native}$) (*parB::cfp-parB)* and inducible *dnaA* ($P_{inducible}$) (*dnaA*::Ω, *parB::cfp-parB, $P_{van}$ dnaA)* grown minimal media (M2G). Super plots showing no significant difference in cell width analysis of mixed population. Small dots represent data points from three independent replicates, large dots represent median values (blue, pink, yellow). The horizontal line represents the mean of three median values. **B.** *dnaA* induced from $P_{xyl}$ (*dnaA*::Ω, *parB::cfp-parB,* $P_{xyl}$) shows ~20% decrease in cell size when grown in minimal media in comparison to WT ($P_{native}$) (*parB::cfp-parB)*. **C.** *dnaA* induced from $P_{xyl}$ (*dnaA*::Ω, *parB::cfp-parB,* $P_{xyl}$) shows no significant difference in cell width grown in M2G in comparison to WT ($P_{native}$) (*parB::cfp-parB).* **D.** Western blot showing DnaA levels of WT (*parB*::CFP-*parB)* and inducible *dnaA* (*dnaA*::Ω, *parB::cfp-parB,* $P_{van}$ *dnaA)*. Cells were grown overnight in minimal media (M2G) to exponential phase with inducer 100μM van before sample preparation. **E.** Quantification of western blot showing significant differences in *dnaA* inducible, 3-fold more DnaA vs WT. Data points show mean ± SD. A parametric t-test was performed using population mean values (N = 3) to compare values for each measurement. p(<0.0001) ****, p(0.0001)***, p(<0.05)*, ns (non-significant). All samples were blinded n= ~600 cells in panel A and ~300 cells in panel B and C.

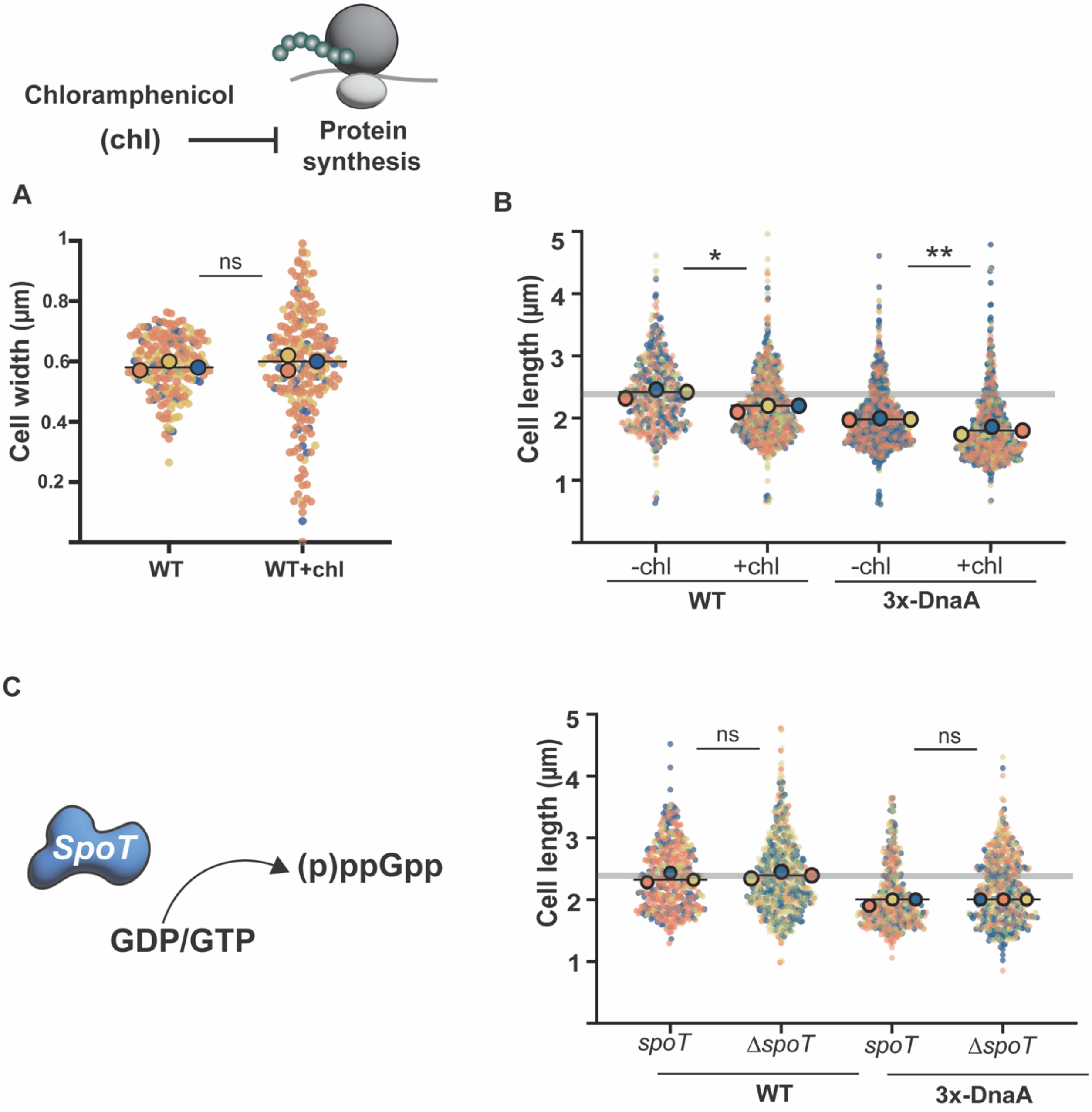


**Figure S2. Impact of protein synthesis A.** inhibition on cell size. WT (*parB*::CFP-*parB)* and inducible *dnaA* (3x-DnaA) (*dnaA*::Ω, *parB::cfp-parB, P*$_{van}$ *dnaA)* cells were grown overnight in minimal media (M2G) overnight with sub-lethal concentration of chloramphenicol and inducer 100µM van. Super plots showing cell length analysis of mixed population. Small dots represent data points from three independent replicates, large dots represent median values (blue,pink,yellow). WT and 3x-DnaA cells show no significant difference in cell width. **B.** Significant differences in cell length were observed. WT cell length reduced cell length approximately 10%, 3x-DnaA cells show a further approximately 10 % reduction when exposed to chloramphenicol. **Impact of ppGpp on**

**cell size. C.** WT delta *spoT (parB*::CFP-*parB* Δ*spoT)* in comparison to 3x-DnaA delta *spoT* (*dnaA*::Ω, *parB::cfp-parB,* $P_{van}$ *dnaA* Δ*spoT*). Super plots showing cell length analysis of mixed population grown in M2G, grown overnight. Cell length analysis shows non-significant changes between 3x-DnaA and 3x-DnaA delta *spoT*, shows similar decrease in cell length ~20%. Data points show mean ± SD. A parametric t test was performed using population mean values (N = 3). p(<0.0001) ****, p(0.0001)***, p(<0.05)*, ns (non-significant). All samples were blinded n= ~600 cells.

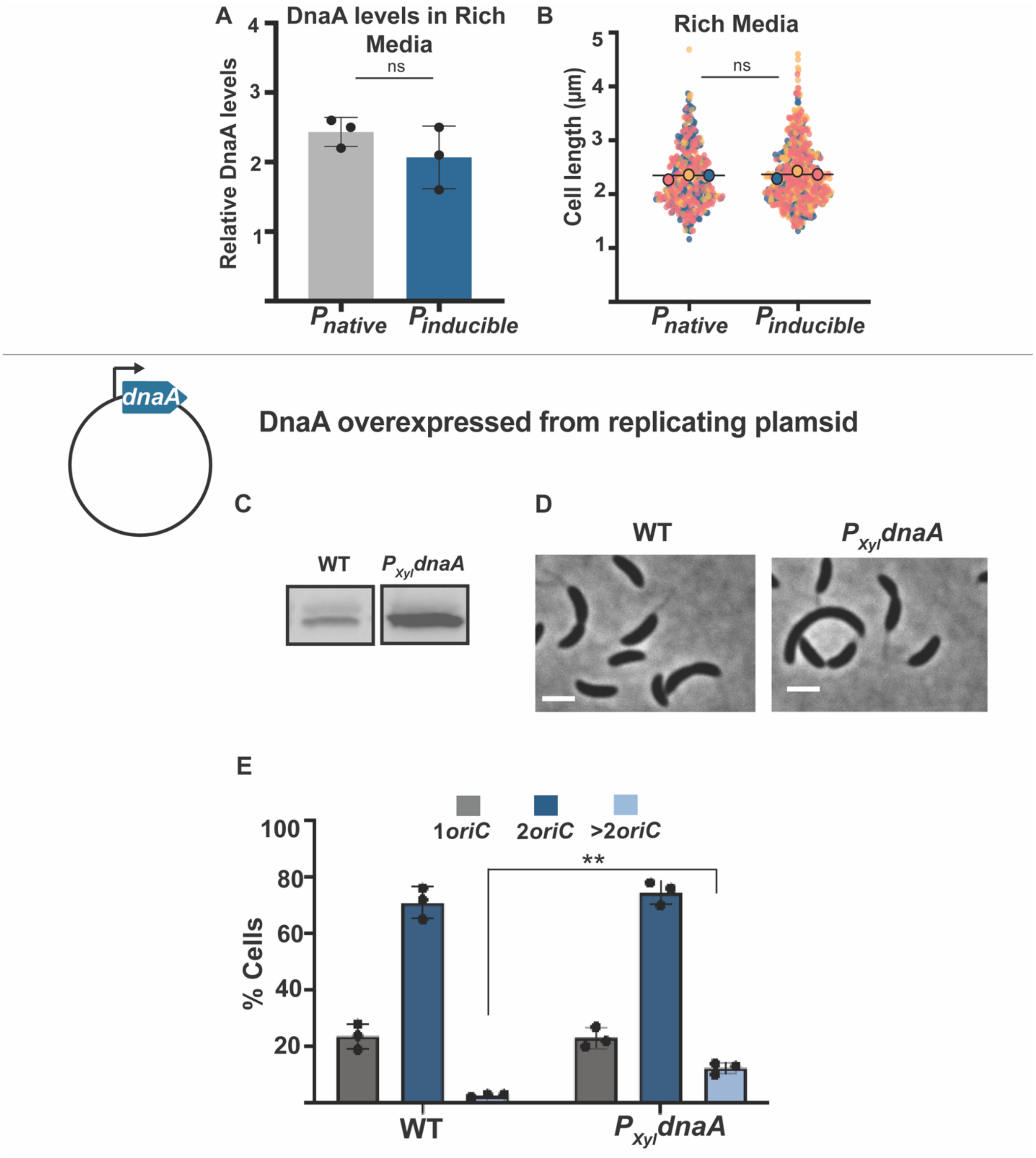


**Figure S3. Nutrient availability and cell size**. **A.** Quantification of western blot showing non-significant differences between WT(*parB*::CFP-*parB)* and inducible *dnaA* (*dnaA*::Ω, *parB::cfp-parB, P*$_{van}$ *dnaA)* in PYE. Cells were grown overnight in rich media (PYE) to exponential phase with inducer 100µM van before sample preparation. **B.** Super plots showing cell length analysis of mixed population grown in PYE. *dnaA* inducible strain show no significant decrease in cell length in comparison to WT. Small dots represent data points from three independent replicates, large dots represent median values (blue, pink, yellow). The horizontal line represents the mean of three median values. n=~600 cells. A parametric t test was performed using population mean values (N = 3). p(<0.0001) ****, p(0.0001)***, p(<0.05)*, ns (non-significant). **C.** Western blot showing DnaA levels of WT and inducible *dnaA* from high copy plasmid (pBXMCS $P_{xyl}$ *dnaA*) in rich media. Cells were grown overnight in minimal media (M2G) to exponential phase with inducer 0.3% xylose before sample preparation **D.** Phase contrast images showing WT and (pBXMCS $P_{xyl}$ *dnaA)* showing over initiation of replication multiple *oriC*, scale 2μm. **E.** Quantification of *oriC, dnaA* induced from a high copy plasmid shows a population (13% cells) that over-initiated replication (>2ori). Data are mean averages from three independent trials. Error bars are ±SEM with two-way ANOVA analysis. p(<0.0001) ****, p(0.0001)***, p(<0.05)*, ns (non-significant).

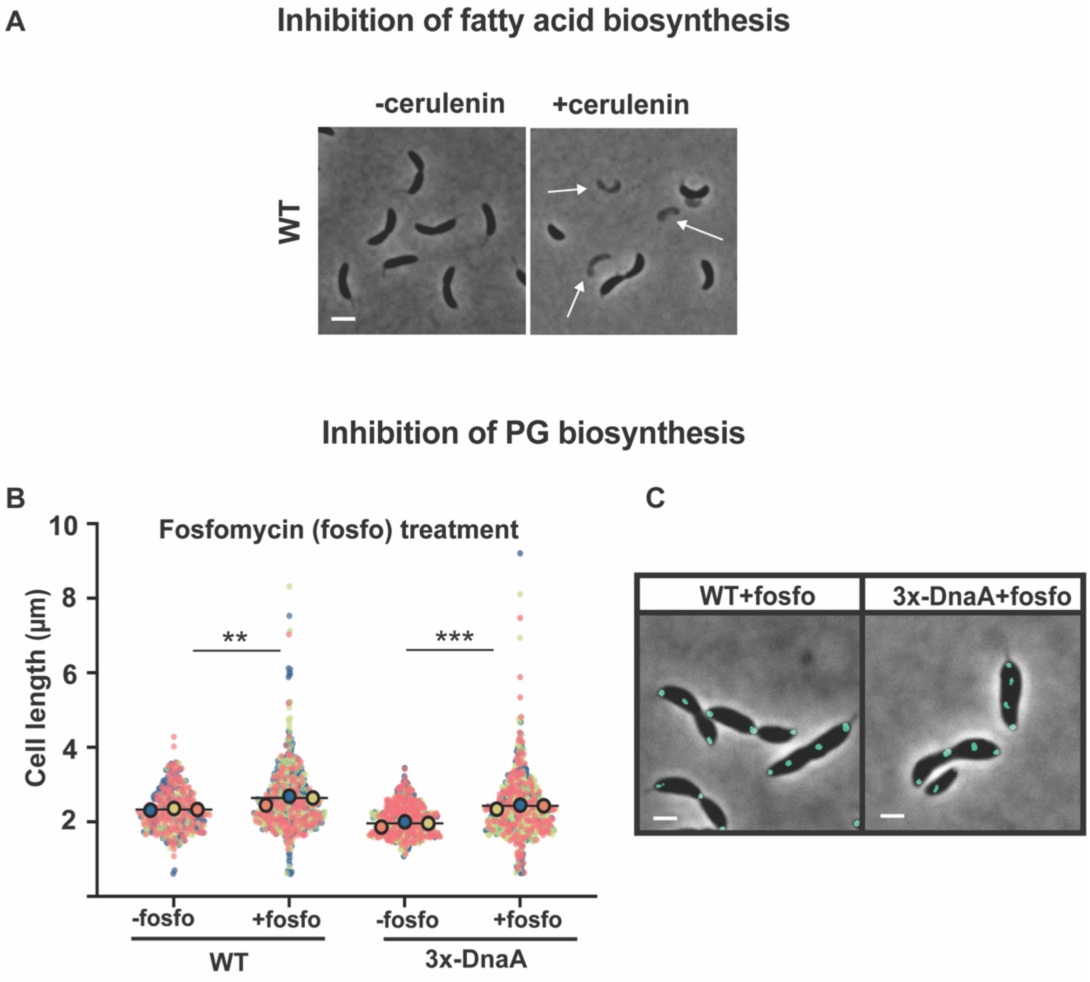


**Figure S4. Envelope biosynthesis and cell size. A** WT (*parB*::CFP-*parB)* cells were grown overnight in sublethal concentration. Representative phase contrast images show that cells were sensitive to 0.6µg of cerulenin, dead cells are highlighted with arrows, scale 2µm. **B.** WT(*parB*::CFP-*parB)* and inducible *dnaA* (3x-DnaA) (*dnaA*::Ω, *parB::cfp-parB, $P_{van}$ dnaA)* in the presence and absence of 5µg Fosfomycin and inducer 100µM van. Super plots showing cell length analysis of mixed population. Small dots represent data points from three independent replicates, large dots represent median values (blue,pink,yellow). **C.** Phase contrast microscopy shows over initiation of replication, CFP-ParB was used to track *oriC*, scale 2µm. A parametric t test was performed using population mean values (N = 3) to compare values for each measurement. p(<0.0001) ****, p(0.0001)***, p(<0.05)*, ns (non-significant). n= ~500 cells.

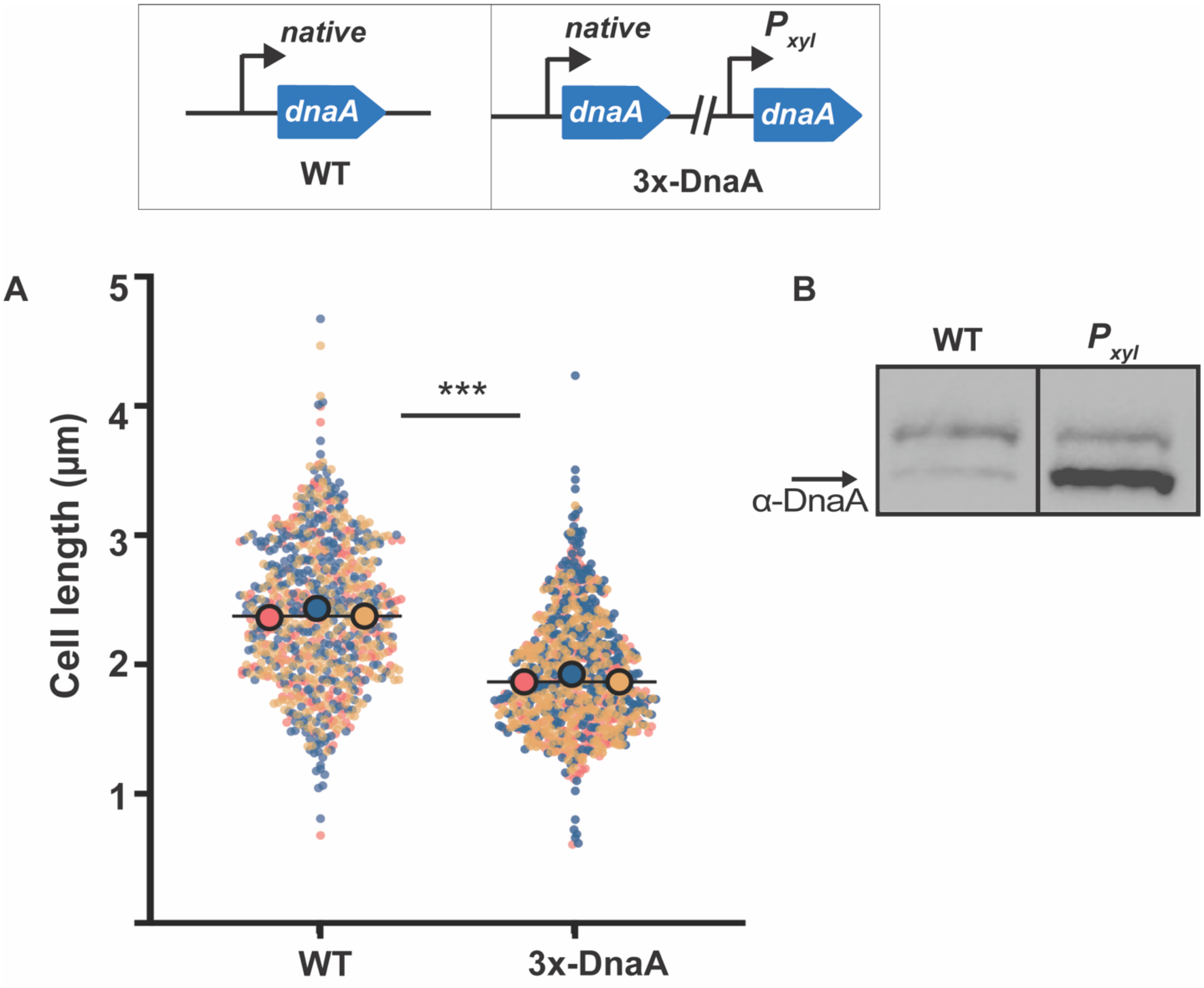


**Figure S5 A. *dnaA* merodiploid cell length. A.** WT (*parB::cfp-parB)* and *dnaA* merodiploid (*parB::cfp-parB, $P_{xyl}$ dnaA)* (3x-DnaA) grown minimal media M2X (xylose for induction). Super plots showing significant difference in cell length analysis of mixed population. Small dots represent data points from three independent replicates, large dots represent median values (blue, pink, yellow). The horizontal line represents the mean of three median values. *dnaA* merodiploid (3x-DnaA) shows ~20% decrease in cell length in comparison to WT. **B.** Western blot showing 3-fold more DnaA in *dnaA* merodiploid (3x-DnaA) in comparison to WT. A parametric t-test was performed using population mean values (N = 3) to compare values for each measurement. p(<0.0001) ****, p(0.0001)***, p(<0.05)*, ns (non-significant). n=~600 cells.

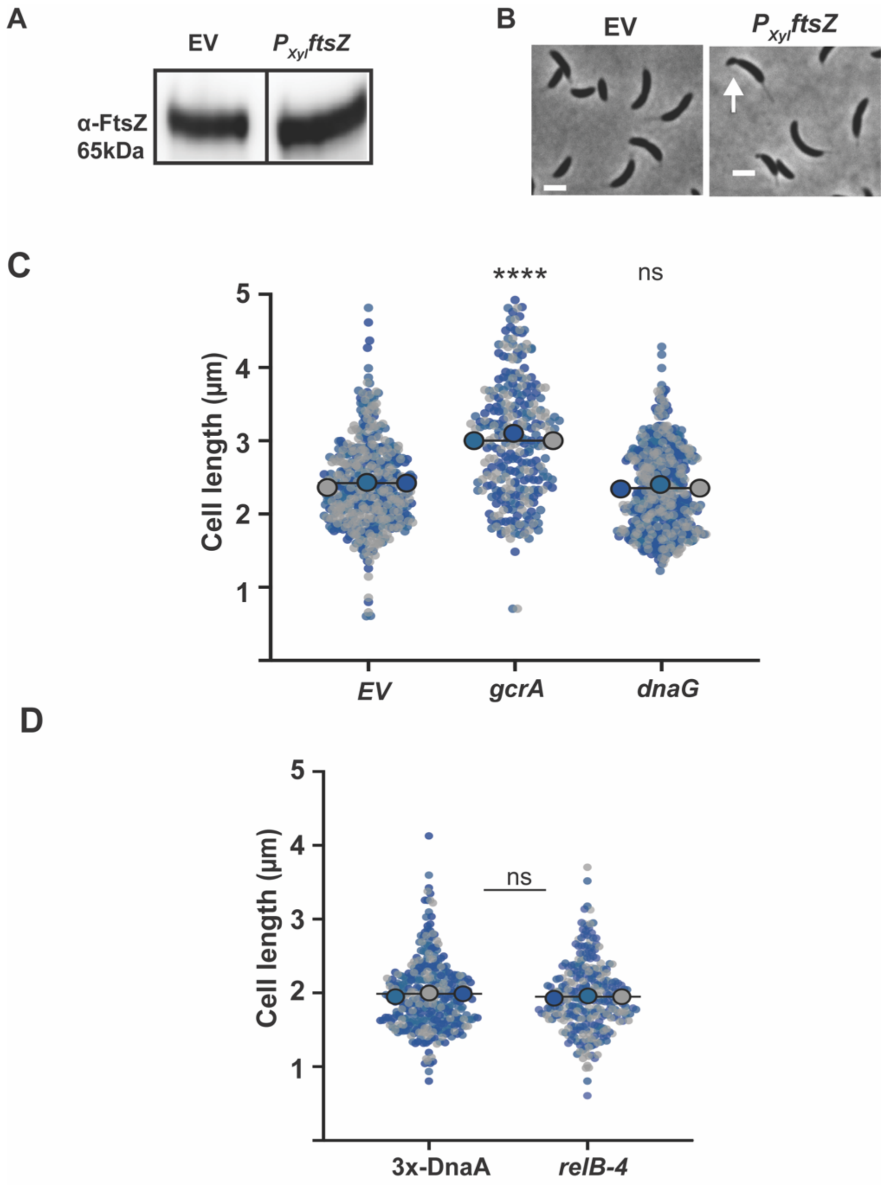


**Figure S6. 3x-DnaA cells display changes in transcriptional profiles. A**. Empty vector (*parB::cfp-parB* $P_{xyl}$ EV) and *ftsZ* merodiploid *(parB::cfp-parB* $P_{xyl}$*-ftsZ)* were grown overnight in M2X (xylose for induction) to exponential phase before sample prepration. Western blot showing FtsZ levels of empty vector (EV) and merodiploid *ftsZ.* **B.** Phase contrast microscopy of EV and *(*$P_{xyl}$*-ftsZ).* Formation of minicells (white arrow) when *ftsZ* is overexpressed shown with arrow. **C.** Super plots representing cell length analysis of *gcrA (parB::cfp-parB* $P_{xyl}$*-gcrA)* and *dnaG* merodiploid *(parB::cfp-parB* $P_{xyl}$*-dnaG)* in comparison to empty vector (*parB::cfp-parB* $P_{xyl}$ EV). Small dots represent data points from three independent replicates, large dots represent median values (blue, navy, grey). Cells were grown overnight in M2X (xylose for induction) to exponential phase.

Overexpression of gcrA increases cell length, whereas dnaG overexpression show non-significant changes. **D.** Super plots representing cell length analysis of *dnaA inducible 3x-DnaA (dnaA*::Ω, *parB::cfp-parB, $P_{van}$ dnaA) and relB-4* merodiploid (*dnaA*::Ω, *parB::cfp-parB, $P_{van}$ dnaA)$P_{xyl}$-relb-4*). Cells were grown overnight in M2X and 100μM van. *relB-4* merodiploid showed non-significant change in comparison to 3x-DnaA. Data show mean ± SD; parametric t-test performed using population mean (N = 3). p(<0.0001) ****, p(0.0001)***, p(<0.05)*, ns (non-significant). All samples blinded, n = ~500 cells.

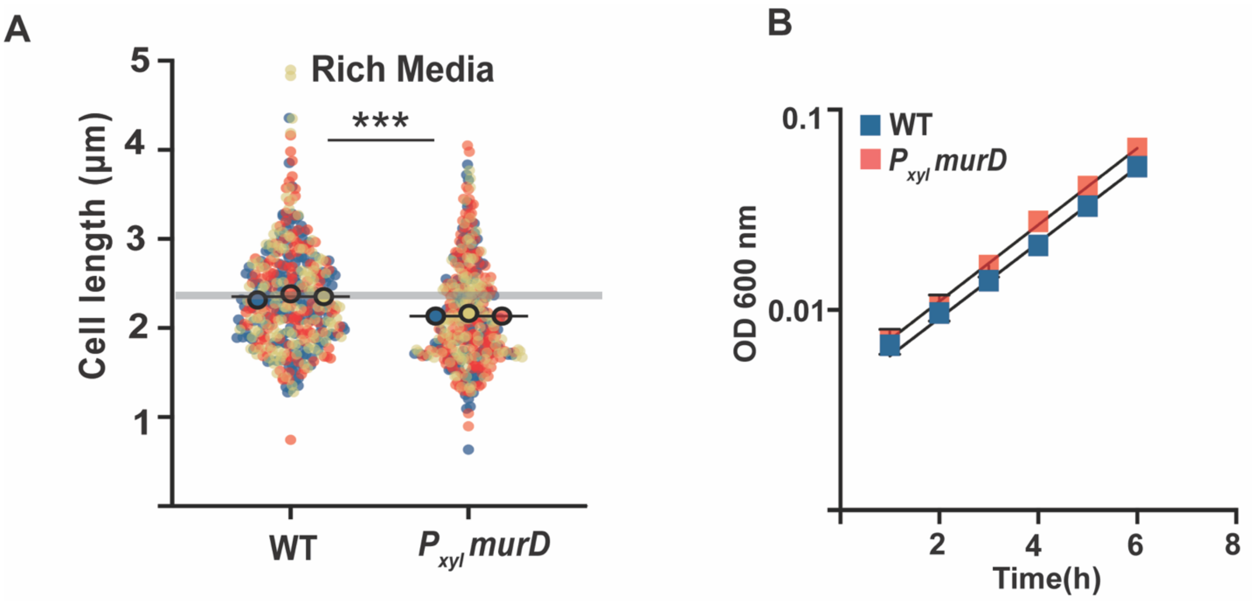


**Figure S7. MurD impact cell size in rich media. A.** Phase contrast images of WT(*parB::cfp-parB) murD* merodiploid *(parB::cfp-parB $P_{xyl}$-murD)*, scale 2µm. **B.** Super plots of cell length analysis show *murD* merodiploid decreases cell length by 10% in comparison to WT. Small dots represent data points from three independent replicates, large dots represent median values (blue, orange, yellow). Cells were grown overnight in in rich media (PYE) and PYEX (xylose 0.2%) to exponential phase. All samples blinded, n = ~550 cells. **C.** Growth curves show doubling time remains same when *murD* is overexpressed. Data show mean ± SD; parametric t-test performed using population mean (N = 3). p(<0.0001) ****, p(0.0001)***, p(<0.05)*, ns (non-significant). All samples blinded, n = ~500 cells.

**Table S1.** RNA-seq data 3x-DnaA compared to EV

| | | EV vs 3x-DnaA | | EV vs 3x-K195I | |
|---|---|---|---|---|---|
| **Gene ID** | **Description** | **FC** | **FDR** | **FC** | **FDR** |
| **CCNA_02793** | **S9 family peptidase** | **3.3** | **1.60E-10** | **NS** | **NS** |
| **CCNA_02792** | **TonB-dependent outer membrane receptor** | **3.1** | **7.27E-12** | **NS** | **NS** |
| **CCNA_03227** | **TonB-dependent receptor** | **1.9** | **1.52E-02** | **NS** | **NS** |
| **CCNA_02328** | **cell cycle sigma 70 cofactor GcrA** | **1.8** | **3.56E-07** | **NS** | **NS** |
| **CCNA_03144** | **DNA primase (*dnaG)*** | **1.8** | **7.54E-07** | **NS** | **NS** |
| **CCNA_02891** | **GanA-family beta-galactosidase** | **1.7** | **1.21E-02** | **NS** | **NS** |
| **CCNA_02976** | **MCP-signal associated domain protein** | **1.7** | **3.46E-02** | **NS** | **NS** |
| **CCNA_02640** | **phospho-N-acetylmuramoyl-pentapeptide-transferase *(mraY)*** | **1.6** | **1.63E-02** | **NS** | **NS** |
| **CCNA_03270** | **AsnC-family transcriptional regulator** | **1.6** | **3.53E-02** | **NS** | **NS** |
| **CCNA_02902** | **xylan alpha-1,2-glucuronosidase** | **1.5** | **1.89E-03** | **NS** | **NS** |
| **CCNA_02623** | **cell division protein FtsZ** | **1.5** | **6.00E-05** | **NS** | **NS** |
| **CCNA_01485** | **aldose 1-epimerase** | **1.5** | **2.18E-02** | **NS** | **NS** |
| **CCNA_02639** | **UDP-N-acetylmuramoylalanine--D-glutamate ligase (*murD*)** | **1.5** | **1.99E-03** | **NS** | **NS** |
| **CCNA_01003** | **flagellar biosynthesis protein FliO** | **1.5** | **3.48E-02** | **NS** | **NS** |
| **CCNA_01979** | **LexA repressor** | **1.5** | **3.46E-02** | **NS** | **NS** |
| **CCNA_01116** | **histidine protein kinase DivJ** | **1.5** | **9.31E-03** | **NS** | **NS** |
| **CCNA_01489** | **alpha-L-arabinofuranosidase** | **1.5** | **3.37E-02** | **NS** | **NS** |
| **CCNA_03105** | **DnaJ domain protein** | **-1.5** | **1.09E-02** | **NS** | **NS** |
| **CCNA_02814** | **transposase** | **-1.5** | **1.52E-02** | **NS** | **NS** |
| **CCNA_03426** | **MarR family repressor protein** | **-1.8** | **1.09E-02** | **NS** | **NS** |
| **CCNA_01247** | **CESA-like glycosyltransferase** | **-2.6** | **2.74E-03** | **NS** | **NS** |
| **CCNA_03231** | **anti-toxin protein relB-4** | **-2.9** | **2.85E-05** | **NS** | **NS** |
| CCNA_03229 | FAD dependent oxidoreductase | 3.4 | 6.50E-06 | 2.6 | 4.30E-04 |
| CCNA_00008 | chromosome replication initiator protein *(dnaA)* | 3.0 | 4.30E-09 | 3.4 | 1.96E-10 |
| CCNA_01783 | chromosome replication regulator protein (*hdaA)* | 2.4 | 5.83E-04 | 2.5 | 5.97E-04 |
| CCNA_03988 | hypothetical protein | 2.4 | 5.08E-04 | 2.0 | 5.52E-03 |
| CCNA_00655 | hypothetical protein | 2.2 | 2.24E-04 | 2.3 | 6.51E-05 |
| CCNA_03481 | DNA integration/recombination/inversion protein | 2.0 | 3.53E-02 | 2.1 | 1.70E-02 |

| | | | | | |
|---|---|---|---|---|---|
| CCNA_01966 | vitamin B12-dependent ribonucleotide reductase (*nrdJ*) | 1.9 | 1.03E-08 | 1.5 | 8.17E-05 |
| CCNA_01780 | Exopolyphosphatase (*ppx1)* | 1.9 | 4.37E-05 | 2.2 | 4.98E-07 |
| CCNA_01535 | single-strand DNA binding protein | 1.8 | 9.65E-05 | 2.0 | 3.4E-06 |
| CCNA_02635 | SEDS-family peptidoglycan polymerase FtsW | 1.8 | 1.07E-03 | 1.5 | 2.78E-02 |
| CCNA_03892 | HD family metal-dependent phosphohydrolase | 1.7 | 1.41E-04 | 1.8 | 9.93E-05 |
| CCNA_02246 | division plane positioning ATPase MipZ | 1.7 | 1.35E-05 | 1.6 | 6.28E-05 |
| CCNA_01791 | CTP synthase | 1.6 | 1.36E-03 | 1.4 | 4.80 E-05 |
| CCNA_02037 | ATP-dependent endopeptidase Lon | 1.6 | 3.53E-02 | 1.7 | 6.34E-03 |
| CCNA_02363 | hypothetical protein | 1.6 | 1.86E-02 | 1.7 | 3.32E-03 |
| CCNA_01517 | NUDIX hydrolase family protein | 1.6 | 1.77E-02 | 1.7 | 3.82E-03 |
| CCNA_01486 | GguC-family protein | 1.5 | 1.93E-02 | 1.5 | 1.93E-02 |
| CCNA_02087 | deoxyguanosinetriphosphate triphosphohydrolase | -1.5 | 7.39E-05 | -1.8 | 1.38E-08 |
| CCNA_00218 | conserved hypothetical protein | -1.6 | 8.18E-03 | -1.6 | 1.3E-02 |
| CCNA_01115 | two component sensor histidine kinase CckN | -1.7 | 7.55E-03 | -1.6 | 3.2E-02 |
| CCNA_01606 | Soj/ParA-related ATPase protein | -2.2 | 2.10E-02 | -3.1 | 4.34E-04 |
| CCNA_02872 | phage phi-C31 gp36 major capsid-like protein | -2.7 | 5.32E-04 | -1.9 | 4.34E-02 |

Column showing RNA-seq data EV vs 3x-DnaA and EV vs K195I. Highlighted genes were only upregulated in 3xDnaA vs K195I. Non-highlighted were upregulated significantly in both 3x-DnaA and K195I. Fold change (FC) ≥1.5, False discovery rate value (FDR) ≤ 0.05

NS= non-significant change

**Table S2.** List of strains used in this study.

| Name | Relevant genotype | Reference |
|---|---|---|
| PM293 | CB15N, *parB*::cfp-*parB* | [23] |
| PM109 | CB15N, *parB*::cfp-*parB ΔvanA* $P_{van}$-*dnaA*-aaCI *dnaA*::Ω (spec/strep) $spec^R$,$strep^R$ | [23] |
| PM138 | CB15N Δ*vanA parB*::CFP-*parB xylX*::*dnaA dnaA*::Ω-(spec/strp) $gent^R$ $spec^R$,$strep^R$ | [23] |
| PM1474 | CB15N, *parB*::cfp-*parB* $P_{xyl}$ *dnaA (pBXMCS-2) (*$kan^R$) | This study, [10] |
| PM672 | CB15N *parB*::CFP-*parB ΔspoT ΔvanA (*$kan^R$) | This study |
| PM806 | CB15N Δ*vanA parB*::CFP-*parB* $P_{van}$-*dnaA* (aaC1) *dnaA*::W-(spec/strp) Δ*spoT* $spec^R$,$strep^R$,$kan^R$ | This study |
| PM1393 | (CB15N *parB*::CFP-*parB dnaN*::*dnaN* mcherry) ($kan^R$) | This study, [24] |
| PM123 | (CB15N Δ*vanA parB*::CFP-*parB vanA*::*dnaA dnaA*::Ω-(spec/strp) *dnaN*::*dnaN* mcherry) ($gent^R$, $spec^R$,$strep^R$,$kan^R$) | This study |
| PM71 | (CB15N CFP-*parB PvanA*-mCherry-*parB*(pMT1) *parS*(pMT1)-*pleC* ($gent^R$,$kan^R$) | This study, [25] |
| PM74 | CB15N CFP-*parB PvanA*-mCherry-*parB*(pMT1) *parS*(pMT1)-$P_{xyl}$-*dnaA*($gent^R$,$kan^R$,$tet^R$) | This study |
| PM566 | CB15N, *parB::cfp-parB,* $P_{xyl}$ *empty-vector(pXCHYC-2)* ($kan^R$) | [26] |
| PM559 | CB15N *parB*::CFP-*parB* $P_{xyl}$ *dnaA* $^{WT}$ ($kan^R$) | This study |
| PM1370 | CB15N *parB*::CFP-*parB* $P_{xyl}$ *dnaA* $^{\Delta ala}$ ($kan^R$) | This study |
| PM875 | CB15N *parB*::CFP-*parB* $P_{xyl}$ *dnaA* $^{K195I}$ ($kan^R$) | This study |
| PM838 | CB15N *parB*::CFP-*parB* $P_{xyl}$ *dnaA* $^{I,II,III\ \Delta IV}$ ($kan^R$) | This study |
| PM776 | CB15N *parB*::CFP-*parB* $P_{xyl}$ *ftsZ* $^{WT}$ ($kan^R$) | This study |
| PM1123 | CB15N *parB*::CFP-*parB* $P_{xyl}$ *gcrA* $^{WT}$ ($kan^R$) | This study |
| PM1480 | CB15N *parB*::CFP-*parB* $P_{xyl}$ *dnaG* $^{WT}$ ($kan^R$) | This study |
| PM1476 | CB15N *parB*::CFP-*parB* $P_{xyl}$ *mraY* $^{WT}$ ($kan^R$) | This study |
| PM1195 | CB15N *parB*::CFP-*parB* $P_{xyl}$ *murD* $^{WT}$ ($kan^R$) | This study |
| PM1472 | CB15N Δ*vanA parB*::CFP-*parB Pvan-dnaA* (aaC1) *dnaA*::W-(spec/strp) $P_{xyl}$ *murD* $^{WT}$ $spec^R$,$strep^R$,$kan^R$ | This study |

**Table S3.** List of plasmids used in this study.

| Name | Relevant genotype | Reference |
|---|---|---|
| pNPTS138 | Nonreplicating vector for allelic replacement (kan$^R$ ) *oriT sacB* | [1] |
| pXCHYC-2 | Integrating constructs encoding C-terminal mCherry fusions under the control of native $P_{xyl}$ (kan$^R$) | [1] |
| pBXMCS-2 | medium-high copy plasmid containing multiple cloning site downstream of the $P_{xyl}$ promoter (kan$^R$) | [1,23] |
| pDNA 555 | *dnaA* $^{WT}$ cloned into pBXMCS-2 | This study, [10] |
| pDNA 293 | pNPTS138 derivative to delete native *spoT* | This study |
| pDNA 30 | *dnaN*-mcherry cloned into pCHYPC-2 | [24] |
| pDNA 2 | *dnaA*$^{WT}$ cloned into pXCHYC-2(Δ-mcherry) | This study |
| pDNA 523 | *dnaA*$^{\Delta ala}$ cloned into pXCHYC-2(Δ-mcherry) | This study |
| pDNA 1 | *dnaA*$^{K195I}$ cloned into pXCHYC-2(Δ-mcherry) | [23] |
| pDNA 86 | *dnaA,* $^{I,II,III}$ cloned into pXCHYC-2(Δ-mcherry) | This study |
| pDNA 333 | *ftsZ* $^{WT}$ cloned into pXCHYC-2 (Δ-mcherry) | This study |
| pDNA 435 | *gcrA* $^{WT}$ cloned into pXCHYC-2 (Δ-mcherry) | This study |
| pDNA 558 | *dnaG* $^{WT}$ cloned into pXCHYC-2 (Δ-mcherry) | This study |
| pDNA 556 | *mraY* $^{WT}$ cloned into pXCHYC-2 (Δ-mcherry) | This study |
| pDNA 469 | *murD* $^{WT}$ cloned into pXCHYC-2 (Δ-mcherry) | This study |
| pDNA 623 | 200bp upstream *mraY* in pXCHYC-2(Δ-mcherry) | This study |
